# SARS-CoV-2 M^pro^ protease variants of concern display altered viral and host target processing but retain potency towards antivirals

**DOI:** 10.1101/2023.01.28.525917

**Authors:** Sizhu Amelia Chen, Elena Arutyunova, Jimmy Lu, Muhammad Bashir Khan, Wioletta Rut, Mikolaj Zmudzinski, Shima Shahbaz, Jegan Iyyathurai, Eman Moussa, Zoe Turner, Bing Bai, Tess Lamer, James A. Nieman, John C. Vederas, Olivier Julien, Marcin Drag, Shokrollah Elahi, Howard S. Young, M. Joanne Lemieux

## Abstract

Main protease of SARS-CoV-2 (M^pro^) is the most promising drug target against coronaviruses due to its essential role in virus replication. With newly emerging variants there is a concern that mutations in M^pro^ may alter structural and functional properties of protease and subsequently the potency of existing and potential antivirals. We explored the effect of 31 mutations belonging to 5 variants of concern (VOC) on catalytic parameters and substrate specificity, which revealed changes in substrate binding and rate of cleavage of a viral peptide. Crystal structures of 11 M^pro^ mutants provided structural insight into their altered functionality. Additionally, we show M^pro^ mutations influence proteolysis of an immunomodulatory host protein Galectin-8 (Gal-8) and subsequent significant decrease in cytokine secretion, providing evidence for alterations in escape of host-antiviral mechanisms. Accordingly, mutations associated with the highly virulent Delta VOC resulted in significant increase in Gal-8 cleavage. Importantly, IC50s of nirmatrelvir (Pfizer) and our irreversible inhibitor AVI-8053 demonstrated no changes in potency for both drugs for all mutants, suggesting M^pro^ will remain a high-priority antiviral drug candidate as SARS-CoV-2 evolves.

## INTRODUCTION

The ongoing COVID-19 pandemic caused by severe acute respiratory syndrome coronavirus 2 (SARS-CoV-2) continues to be a significant threat to global public health (1, 2). SARS-CoV-2 is a positive-sense single-stranded RNA virus with a low genome stability and a high mutation rate (3-5). The emergence and rapid spread of SARS-CoV-2 variants have posed a greater challenge in the control of the pandemic due to increased transmissibility, disease severity or resistance towards vaccines and therapies (6-10). Owing to the potential threats of the variant (11), Alpha (B.1.1.7), Beta (B.1.351), Gamma (P.1), Delta (B.1.617.2), and most recently Omicron (B.1.1.529) have been classified into variants of concerns (VOCs) by the World Health Organization (WHO) (12). The genomic sequences of new variants are generated and submitted on the Global Initiative on Sharing All Influenza Data (GISAID) with an unprecedented speed, which aids in further understanding of epidemiology (13) and the distribution of mutations that may lead to changes in viral characteristics (14).

SARS-CoV-2 genome encodes two overlapping polyproteins, pp1a and pp1ab, that are proteolytically processed to generate 16 nonstructural proteins (Nsps), followed by four structural proteins that include the envelope (E), membrane (M), nucleocapsid (N), and spike proteins at the 3′ terminus of the genome (15, 16). Multiple mutations occur in various regions of the viral genome (17). Mutations in the spike protein, which mediates viral entry into human cells (5), is one primary focus of the current research of SARS-CoV-2 variants due to their high impact on infectivity (18), transmissibility (19) and potential immune escape from antibodies (20). However, limited study has been done to examine the mutations in other essential viral proteins. Notably, single point mutations found in SARS-CoV-2 variants are also observed in nsp5 gene, which encodes the viral main protease (M^pro^, also called 3CLpro) (21). M^pro^ cleaves the polyproteins at 11 positions with a consensus cleavage sequence Leu Gln | (Ser, Ala, Gly) to release Nsps, which are required to assemble the viral replication-transcription complex (22, 23). Previous studies show that the SARS-CoV and SARS-CoV-2 M^pro^ share highly similar three-dimensional structures with a 96% sequence identity (24, 25). SARS-CoV-2 M^pro^ is a cysteine protease that forms a dimer composed of two protomers (23, 25). Each protomer comprises the chymotrypsin and 3C-like peptidase domains I and II (residues 10-99 and 100-182, respectively), with the active site consisting of a Cys145-His41 catalytic dyad located in the cleft between the two antiparallel-β-barrel domains. The α-helical domain III (residues 198-303) linked to domain II by a long loop regulates the dimer formation where dimerization is required for the catalytic activity of M^pro^ to cleave the viral polyproteins.

In addition to the indispensable role in viral replication, SARS-CoV-2 M^pro^ also plays an essential role in escaping antiviral defense by cleaving host cell proteins. During pathogenic infection type I interferon (INF) controls innate and adaptive immune responses and induces host defense mechanism by activating JAK-STAT signaling pathway, leading to interferon-stimulated gene (ISG) responses (26). M^pro^ of different coronaviruses, including porcine deltacoronavirus (27, 28), porcine epidemic diarrhea virus (29), and feline infectious peritonitis virus (30), disrupts INF-induced pathway by cleaving NF-κB essential modulator (NEMO).

Galectins are essential regulators in host adaptive and innate immune responses (31). Importantly, recent studies by Pablos et al reveal that SARS-CoV-2 M^pro^ contributes to escaping host antiviral responses by cleaving host proteins and identified host Galectin-8 (Gal-8) as an M^pro^ substrate (32). Gal-8 consists of two carbohydrate recognition domains (CRDs) joined by a linker peptide, which binds to glycans on damaged lysosomes or phagosomes upon infections and recruits autophagy adaptors (33, 34), such as NDP52(35). Pablos et al showed M^pro^ cleaves Gal-8 at the short linker region, DLQ158↓ST, which dislocates the CRD1 from the CRD2. In the same study it was demonstrated that during SARS-CoV-2 infection, Gal-8 binds to spike glycoprotein, but the Gal-8:NDP52 complex and subsequent autophagy is disrupted upon Gal-8 cleavage by M^pro^ further demonstrating that proteolytic processing of Gal-8 is an important antiviral mechanism to overcome a host defenses allowing SARS-CoV-2 to escape antiviral xenophagy (32).

Given the critical roles of SARS-CoV-2 M^pro^ in viral replication as well as host immune escape, and the potential vaccine resistance developed by SARS-CoV-2 variants (36), M^pro^ is an attractive target for drug design to treat COVID-19. The orally administered drug Paxlovid™, developed by Pfizer, contains an active component nimatrelvir, a peptidomimetic inhibitor of M^pro^ with a nitrile warhead covalently binding the catalytic cysteine in the active site (37), but also requires a co-dosing with ritonavir, which inactivates the major human drug-metabolizing enzyme, CYP3A4, to enhance pharmacokinetics of the M^pro^ inhibitor. PaxlovidTM was authorized by FDA as the first emergency treatment for COVID-19 in December 2021 (38). Since the start of the pandemic, we have also examined diverse inhibitors that target SARS-CoV-2 M^pro^. Our previous study has shown that the feline prodrug, GC376, which is used to treat feline coronavirus infection (39), was a potent inhibitor for SARS-CoV-2 M^pro^ in vivo (23), and its analogs displayed increased drug efficacy and solubility (40). We also developed novel reversible and irreversible inhibitors with nitrile (41) and α-acyloxymethylketone warheads (42), AVI-8059 and AVI-8053, respectively, that had comparable drug efficacy to nimatrelvir.

With new variants arising over time, different M^pro^ mutations may result in changes of protein structure, catalytic activity, and most importantly, the potential therapeutic approach targeting M^pro^ (21, 43). A recent study has shown that the P108S mutation in SARS-CoV-2 M^pro^ induced structural perturbation around the substrate-binding region, which led to lower enzymic activity and reduced disease severity in patients infected with the SARS-CoV-2 sub-lineage (B.1.1.284) (43). As for the more transmissible Omicron variant, it carries a single-point mutation, P132H, in M^pro^. The crystal structure of P132H M^pro^ in complex with GC376 was similar to the wild-type M^pro^, and both GC376 and nimatrelvir remain potent against P132H M^pro^ (44). However, additional mutations can still occur as SARS-CoV-2 continues to evolve, potentially influencing the M^pro^ structures, leading to the development of drug resistance to M^pro^ inhibitors.

In this study, we examine the effect of 31 point mutations in nsp5 that occurred from Alpha to more recent Omicron VOCs. We show these mutations influence M^pro^ activity, with some enhancing and others decreasing catalytic efficiency of the protease. Furthermore, substrate specificity is also altered for some mutants. We also assess the influence of M^pro^ mutations on cleavage of host substrate Gal-8 and demonstrate that while cleaved Gal-8 is still able to induce the secretion of TNF /IL6 cytokines the levels are significantly reduced. Importantly, two inhibitors - nimatrelvir and our novel M^pro^ inhibitor - remain potent against all variants of SARS-CoV-2 M^pro^. Overall, this study describes important changes in nsp5 over the evolution of the virus that have implications in clinical manifestations.

## RESULTS

### Prevalence of mutations in Nsp5 in SARS-CoV-2 variants

We utilized GISAID Initiative EpiCoV database (https://www.gisaid.org/) to identify and monitor the single point mutations in the nsp5 gene from clinical isolates of different SARS-CoV-2 lineages. The genomes of 5 VOCs were analysed with hCoV-19/Wuhan/WIV04/2019 (WIV04) sequence as an official reference sequence. We selected 31 mutations based on frequency of occurrence as well as the importance for functionality based on location within the protein molecule (Figure 1A, B and Table S1) and grouped them in several hot spots (Figure 1A) - clustered near the active site (L50F, E47K, E47N and S46F), at the end of a beta-sheet behind the active site (D92G, K90R and L98F) and at the dimer interface (G283S, S284G and A285T).

**Figure 1.**
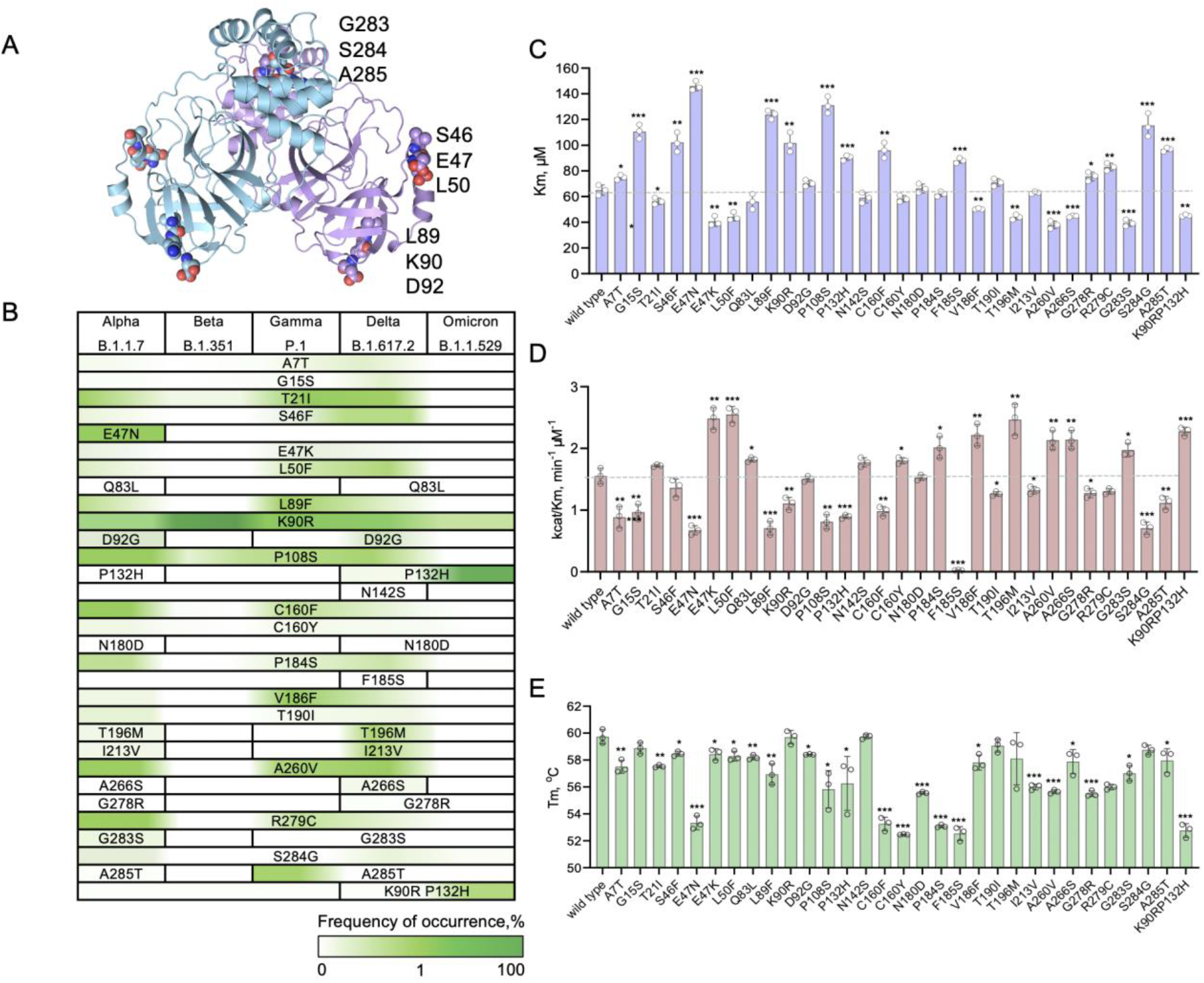
The distribution, prevalence and functional differences of mutations in SARS-CoV-2 M^pro^. **(A)** The crystal structure of SARS-CoV-2 Mpro dimer (PDB 6WTM) mapping VOC mutations from the GISAID database (https://gisaid.org). **(B)** VOC mutation prevalence was calculated and presented as a heat map. **(C)** SARS-CoV-2 M^pro^ mutants reveal differences in K_M_, **(D)** catalytic efficiency (k_cat_/K_M_) and **(E)** Tm values. *: p<0.05 between wild type and the mutants.**: p<0.01. ***: p<0.001. NS: p>0.05.

Two of the most prevalent mutations in NSP5 are K90R and P132H (Figure 1B and Table S1). K90R appeared in Alpha VOC with a prevalence of 23.6% and then became a predominant mutation in Beta VOC (99.9%), with lower prevalence in Gamma and Delta VOCs, 16.2% and 21.7% respectively. The frequency of the K90R mutation became very low in Omicron VOC (0.5%) while the P132H mutation became the most frequent in Omicron VOC (99.9%). Other mutations with relatively high occurrence worth noting are E47N, P108S, C160F, A260V (8.7%, 5.3%, 4.1% and 3.7% in Alpha VOC).

We also chose a double mutation, K90R P132H, which appeared in Delta variant with a very low frequency (0.004%) but became more prevalent in Omicron (0.8%).

### Kinetic parameters and structural alterations in M^pro^ mutants found in five variants of concern

The most significant effect of mutations on proteolytic efficiency of M^pro^ was observed for the residues belonging to Domain I and located near the active site.

L50F is part of the “active site gateway”-a region comprised of two loops L50-Y54 and D187-A191. The mutant exhibited lower KM value, slightly increased kcat and subsequently 1.6-fold higher catalytic efficiency compared to the wild type (Figure 1C, D). The L50F crystal structure revealed loss of charge around S2 binding site and changes in the orientation of side chains of M49 and R188 residues involved in S2 formation. S2 site is known to undergo slight structural changes upon binding to Leu in P2 position (45, 46) and alterations in M49 and R188 side chains may make it more accessible for the substrate, decreasing the KM value (Figure 3A, B insert, Figure S2A, B).

Two substitutions found for E47 residue (E47N and E47K) had drastically different effects on protease activity – polar Asn had less tolerance, leading to a two-fold increase in KM, whereas positively charged Lys had the opposite effect on substrate binding, making the enzyme more efficient (Figure 1C, D). Crystal structure of E47N mutant revealed the change in side chain orientation of amino acids M49, M165, E166 and Q189 involved in S2, S3 and S4 substrate binding sites resulting in the altering surface charge distribution. Also the side chain of Q189 formed weak hydrogen bonds with the backbones of R188 and E166 residues, which are absent in the wild type structure and could affect the formation of S2, S3 and S4 subsites and the substrate binding ability (Figure 3C insert, Figure S2C). Previous crystal structures of M^pro^ with peptidomimetic inhibitors demonstrated that R188 and E166 residues play significant roles in substrate binding by forming hydrogen bonds with Leu in P2 position and a residue in P3 position respectively (23, 24). However, in the E47N mutant, these interactions may not be possible explaining the significant increase in KM value. In the structure of E47K mutant, S2, S3 and S4 binding sites were not altered, but we see that backbone of K47 forms a strong hydrogen bond with the side chain of S46, which is not found in the wild type structure (Figure 3D insert). S46 is a flexible residue and its stabilization aids in opening of the active site making it more accessible. Another feature of E47K mutant worth noting is the drastic change in the charge around S3’ binding site (Figure 3D, Figure S2D).

Charge alteration around S3’ binding site was also observed in the crystal structure of L89F mutant resulting in more hydrophobic surface (Figure 3E, Figure S2E). Besides, Phe and A285T) is the dimer interface (Figure 1A). Despite the fact that A7 residue is part of N-terminus responsible for interaction with C-terminal domain of second monomer (25), and threonine substitution changed its hydrophobicity, we observe only a mild effect of this mutation on activity of protease, reducing the efficiency 1.5 substitution at position 89 enabled pi-stacking interactions with F66 and hydrophobic interactions with V20, which are not observed in the structure of M^pro^ wild type. These changes may affect the flexibility of S3’ substrate binding pocket (Figure 3E insert), which increased the KM value of L89F mutant 2-fold making the enzyme less efficient (Figure 1C, D).

Another region important for M^pro^ functionality, where a cluster of mutations was identified (A7T, G283S, S2684G times (Figure 1C, D). The crystal structure also revealed no significant changes in the area of the active site, except for the orientation of side chains of N142 rotating towards the catalytic dyad as well as M49 and R188, the residues important for the interaction with Leu in P2 position and making S2 site less charged (Figure S2F).

C-terminal region mutations involved in dimerization - S284G and A285T - decreased the catalytic efficiency 2 and 1.4 times respectively, whereas G283S substitution was well tolerated (Figure 1C, D). The crystal structures of G283S and S284G did not result in any interesting changes with the exception of small charge alteration around S2 site (Figure S2G, H).

The mutants with high prevalence – K90R (99.8% in Beta variant), T21I (9.6% in Gamma) or P132H (99.9% in Omicron) resulted in no changes in catalytic efficiency with the exclusion of V186F (7.2% in Gamma variant) and A260V (3.7% in Alpha and 5% in Delta variants) - these mutations rendered protease 1.5 times more efficient (Figure 1C, D). It is worth noting that for most M^pro^ variants the affected parameter was KM suggesting that mutations altered the substrate binding rather than catalysis itself. Crystal structures of K90R, P132H, T190I and A260V also showed no significant changes, which is consistent with the previously reported structure of Omicron M^pro^ (Figure S3I, J, K, L). Interestingly, the catalytic efficiency for the double mutant K90, P132H showed a higher activity revealing a synergistic effect.

Of notable interest was a rare F185S mutation identified in a clinical isolate that had a drastic effect on activity with 30-fold decrease of turnover rate (Figure 1C, D). F185 is located at the base of the D187-A191 loop, which is a part of the “active site gateway” region. The residues on this loop are involved in forming S2 and S4 substrate binding pockets. F185 forms several interactions, which hold the loop together - pi stacking interaction with P184, and hydrophobic interactions with A194 and with A173, which are parts of the beta-sheet close to the active site. All interactions F185 is involved in are at the same plane and aid in stabilizing the loop (Figure S3B). B-factors of that region confirm that the base of the loop where F185 is located is very rigid whereas the tip is flexible. One can predict that Ser substitution would result in a loss of important hydrophobic interactions and destabilization of that region. Size exclusion chromatography of SARS-CoV-2 M^pro^ F185S revealed a shift in retention volume with calculated MW of the protein of 49 kDa suggesting that the mutant exists in a monomeric form but based on Stokes radius it may have an altered shape of the molecule (Figure S4).

### Differential scanning fluorimetry (DSF) reveals the differences in stability of M^pro^ molecule variants

M^pro^ is a stable protein with a Tm of 59.7 °Cand most mutations had a moderate destabilizing effect on protease structure (Figure 1E).

The most vulnerable region in the protease molecule where mutations had a significant destabilizing effect was a region in Domain II. We observed a -7 °Cshift for C160F, C160Y and P184S, -4 °Cfor N180D, -7 °Cfor F185S and -2 °Cfor V186F. C160 is located on a beta sheet and forms a hydrophobic interaction with F112, which is located on the antiparallel beta sheet. B factors derived from the structure of the wild type indicate that the whole region is very rigid, and one can predict that any perturbations in that area would cause destabilization in protein molecule (Figure S3C). The mutations involved in dimerization - A7T, G283S, S284G and A285T – exhibited 2 degrees lower Tm.

The loss of proline at position 108 and 132 affected the stability of the protein molecule as well and resulted in more significant Tm changes (−3.9 °Cfor P108S and -3.5 °Cfor P132H). Interestingly, we also observe significant decrease of Tm for double mutant, K90R P132H (−6.9 °C), whereas the single mutations did not cause changes in thermal stability.

### M^pro^ inhibitors - nirmatrelvir and AVI-8053 - remain potent against the variants

Two peptidomimetic inhibitors against M^pro^ variants - nirmatrelvir (PF-07321332), a component of Paxlovid™ (Pfizer) and AVI-8053, an irreversible inhibitor with an acyloxymethylketone (AMK) warhead (42), developed by Li Ka Shing Applied Virology Institute at the University of Alberta were tested against 31 M^pro^ mutants. We performed the inhibitory studies and confirmed that all variants were inhibited by both drugs with nanomolar IC50 values (Figure 2F), confirming the potency of existing antivirals against VOCs in vitro.

**Figure 2.**
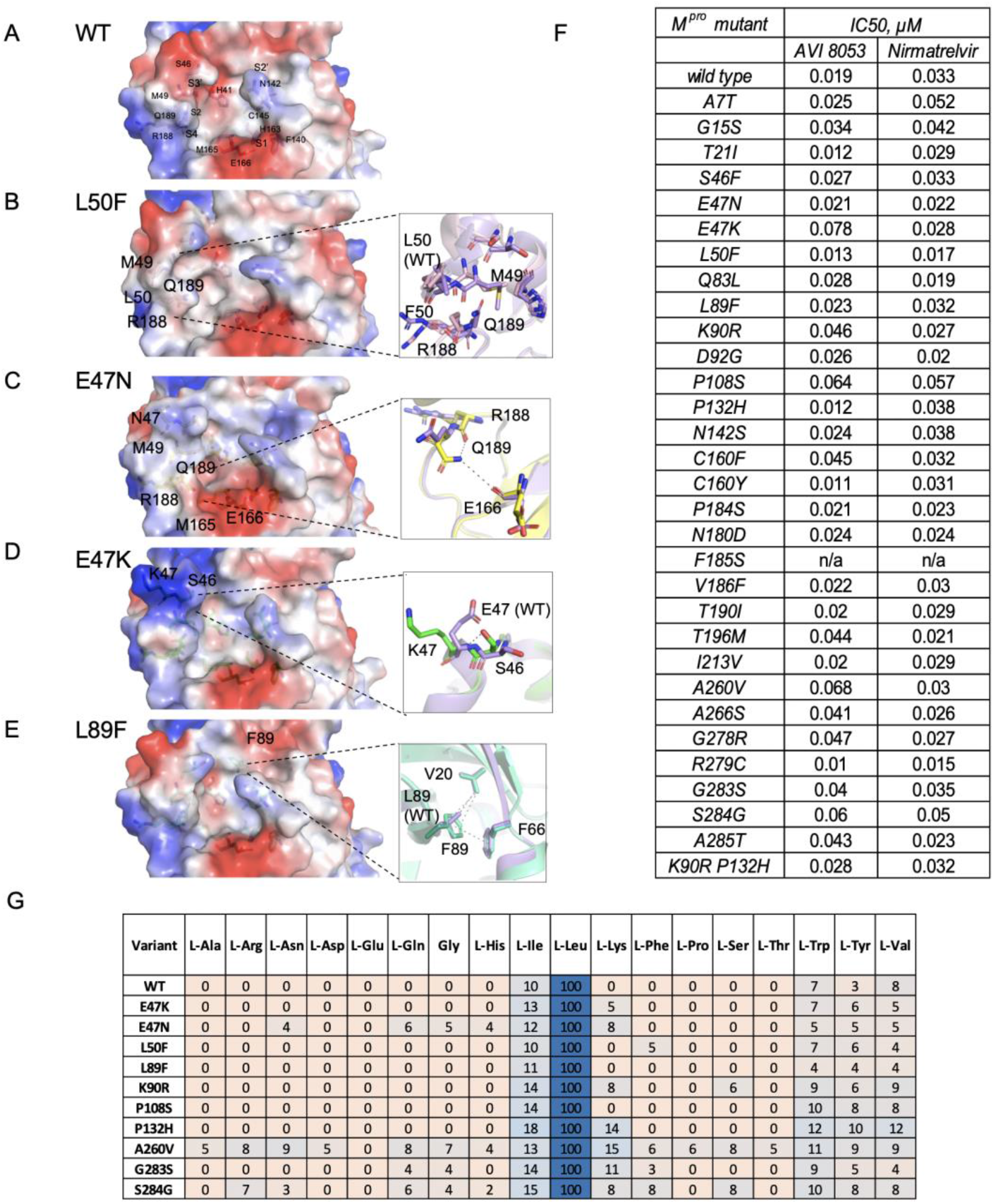
Molecular surface representations showing the electrostatic surface potentials of M^pro^ wild type **(A)**and the mutants **(B-E)**. Hydrogen bonds and hydrophobic interactions between residues of interest are depicted in insets. Wild type structure is indicated by purple. **(F)** IC_50_ values of M^pro^ wild type and the mutants for AVI inhibitor AVI-8053 and nirmatrelvir. **(G)** Substrate specificity profile of SARS-CoV-2 M^pro^ at presented as heat maps using substrate library with natural amino acids assessing the P2 position with Ac-Mix-Mix-P2-Gln-ACC.

### Mutations in M^pro^ change protease substrate specificity

The crystal structures of several variants revealed the changes in side chains orientations of residues in the active site comprising substrate binding pockets, especially S2 and S4, which might lead to altered amino acids preferences. Accordingly, changes in KM values for most of variants were observed (Figure 1C), indicating alterations in a mode of substrate binding.

To reveal if mutations caused changes in M^pro^ substrate preferences we chose 10 mutants with the most significant differences in KM values and deployed Hybrid Combinatorial Substrate Library (HyCoSuL) approach (47). The library consisted of three sublibraries (P2, P3 and P4) of tetrapeptides with two fixed positions, one of each was glutamine at P1 position, as a key feature of SARS-CoV and SARS-CoV-2 main proteases is their ability to cleave the peptide bond after Gln, and two varied positions containing an equimolar mixture of 19 amino acids. P2, P3 or P4 positions in 3 sublibraries correspondingly were represented by 19 natural and over 100 unnatural amino acids. At the C-terminal, an ACC (7-amino-4-carbamoylmethylcoumarin) fluorescent tag was attached at the P1′ position to monitor the cleavage reaction.

The library screen revealed that some variants gained or lost the ability to accommodate certain amino acids in substrate binding pockets (Figure 2G, Figure S5). As it was demonstrated before Leu is the most preferred amino acid at P2 position with very low (≤10%) activity for other mostly hydrophobic natural amino acids, such as L-Ile and L-Val. However, we observed that Omicron (P132H) variant as well as variants with mutations close or at dimer interface (A260V, G283S and S284G) permitted charged L-Lys and had an increased activity towards L-Trp and L-Tyr (Figure 2G). Among all mutants A260V was the most promiscuous at this position – allowing almost all amino acids, even L-Pro with low activity though, which is a striking difference compared to the wild type. We observed the same trend with A260V for unnatural amino acids with this variant becoming the most permissive in comparison to the wild type (Figure S6).

The specificity for P3 position did not change significantly for most of the variants (Figure S5). All of them as well as the wild type preferred hydrophobic and positively charged amino acids – with L-Lys, L-Arg, L-Val and L-Thr having the highest activity. Interestingly, many of the variants lost the ability to cleave the substrate with L-Gln and L-His at P3 position. As described previously, the S3 pocket is not well-defined in the M^pro^ molecule leading to a broad substrate specificity profile for both natural and unnatural amino acids.

P4 position is permissive as well with a highest preference for L-Ala and L-Val for the wild type (Figure S5). Most of the variants resulted in either the same or lower activity and some lost the ability to cleave certain amino acids, like L-Ile, L-Pro, and the non-natural amino acids D-Phg, β-Ala, and L-Agb.

### M^pro^ cleaves host protein Gal-8 at multiple sites

M^pro^ plays an important role in viral infection not only because of its essential function for viral replication but also because of interacting directly with host proteins, as it has been reported recently (32). We sought to study the cleavage of Gal-8, one of M^pro^ host substrate, in more detail and reveal how mutations in viral protease affect its processing.

Gal-8 was recombinantly expressed in E. coli, purified, incubated with M^pro^ and the cleavage was detected by the end point SDS-PAGE-based assay and mass spectrometry (Figure S7). The time course of M^pro^ cleavage of Gal-8 revealed that the proteolysis occurs at several sites at the same time with some sites being more accessible than others and some cleavage products being transient (Figure 3A, B). Mass spectrometry analysis identified three sites of M^pro^ scission - NLQ9↓NI, DLQ158↓ST and FLQ246↓ES, where the DLQ158↓ST site (Figure S7), that was identified previously (32), is located in the linker region between two carbohydrate recognition domains (Figure 3C). We observed that M^pro^ first cuts Gal-8 at Q9 and Q158 sites, producing Gal-8^10-317^ and Gal-8^159-317^ cleavage products (34.8 and 17.9 kDa bands respectively). However, after 6 hours Gal-8 10-317 band disappears after being cut again at Q158 site resulting in Gal-8^10-158^ product (16.8 kDa). The band at 21 kDa represents the intermediate product of cleavage events at Q246, which then was cut further possibly at Q158, resulting in smaller fragments. Thus, incubation of M^pro^ with Gal-8 for 24 hours produced two stable cleavage products: Gal-8^159-317^ (17.9kDa) and Gal-8^10-158^ (16.8kDa).

**Figure 3.**
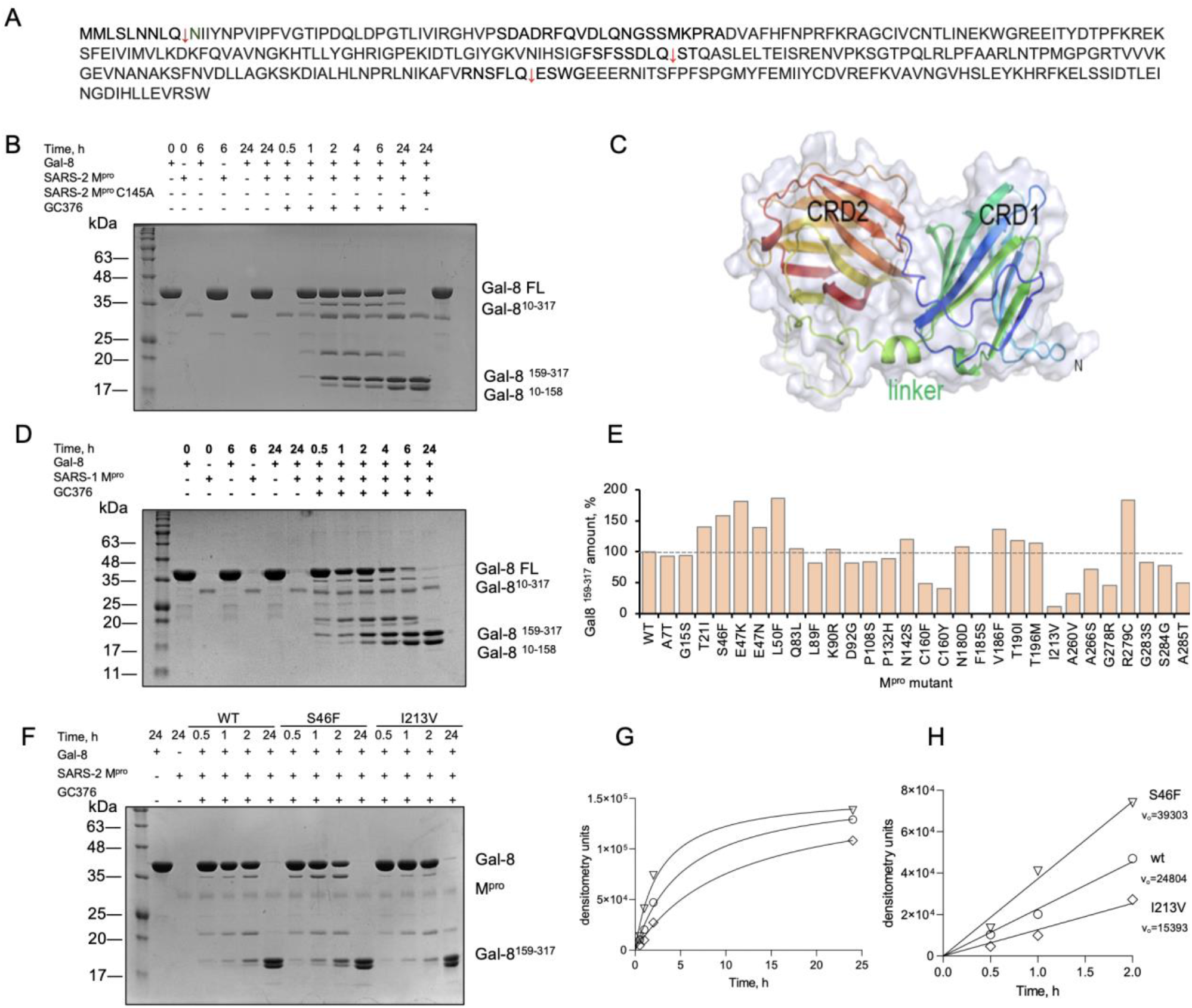
The cleavage of host cell substrate Gal-8 by M^pro^ from SARS-CoV-2 variants and SARS-CoV-1. **(A)** The sequence of Gal-8. The arrows indicate the cleavage sites for viral proteases. **(B)** SDS-PAGE-based time-course of Gal-8 cleavage by SARS-CoV-2 M^pro^. Gal8 was incubated with the protease at 37°C and reaction was stopped at specific time points with M^pro^ inhibitor, GC376. **(C)** Structural model of Gal-8. CRD: carbohydrate recognition domain. **(D)** The time course of Gal-8 cleavage by SARS-CoV-1. **(E)** Gal-8 was cleaved by wild type and M^pro^ mutants. Densitometry analysis of the of generated Gal-8 ^159-317^ product in percent in comparison to Gal-8 cleavage by the wild type. (F) SDS-PAGE gel of the time course of Gal-8 cleavage by M^pro^ wild type, M^pro^ S46F and M^pro^ I213V. **(G)** The dependence of Gal-8^159-317^ band intensities represented in densitometry units on time. **(H)** The linear part of graph B was used to calculate the initial velocities (v_0_) of cleavage reactions.

Since SARS-CoV-2 M^pro^ shares 96% identity in amino acid sequence with SARS-CoV M^pro^ and both enzymes have similar substrate preferences, we were interested to assess if Gal-8 was also a substrate for SARS-CoV M^pro^. We tested the cleavage of Gal-8 by SARS-CoV M^pro^ using the same approach as for SARS-CoV-2 protease and revealed that the cleavage pattern and the rate of proteolysis were very similar (Figure 3D).

Gal-8 belongs to the tandem-repeat-type subclass of the galectin family, same as galectin-9 (Gal-9), which also consists of two carbohydrate recognition domains joined by a linker peptide. The sequence of Gal-9 has two distinct recognition motifs for M^pro^ – Gln in P1 and Leu in P2 positions. Therefore, it was logical to assume that Gal-9 might also be a substrate for M^pro^ protease. To test this hypothesis, Gal-9 was recombinantly expressed in E. coli, purified, and M^pro^ cleavage assay was performed under the same conditions as for Gal-8. However, we did not detect any cleavage products after 24 hours of proteolytic reaction, indicating that Gal-9 is not a substrate of SARS-CoV-2 M^pro^ (Figure S8).

### The cleavage of Gal-8 by M^pro^ variants

The time course of Gal-8 proteolysis by M^pro^ demonstrated that Q158 was the most accessible cleavage site, and Gal-8159-317 product was detectable after 30 min of cleavage and remained stable for at least 24 hours. The dependence of Gal-8159-317 band formation on time, assessed by densitometry, showed a linear relationship within 2 hours (Figure S9A). Therefore, to compare the proteolytic efficiency of variants the intensities of Gal-8^159-317^ bands after 2 hours of Gal-8 cleavage by mutants were assessed by densitometry analysis.

The most significant increase of Gal-8 cleavage rate was observed for the variants with mutations around the active site – S46F, E47K, E47N, L50F, N142S, V186F and R279C; the lowest activity was demonstrated by I213V mutant and also F185S, consistent with the previous observations (Figure 3E, Figure S9C). The more detailed time course analysis of two mutants with highest (S46F) and lowest (I213V) activity confirmed the differences in cleavage rates resulting in 1.5 times increase for S46F and 1.6 times decrease for I213V (Figure 3G and H). The mutations that affected the protein stability (C160F, C160Y and F185A) demonstrated significant decrease in activity, highlighting the importance of that region for M^pro^ functionality.

The cleavage of Gal-8 by M^pro^ may lead to several consequences, one of which is the alteration or loss of Gal-8 functionality. Pablos et al. in a recent study demonstrated that the cleavage event at Q158 resulted in a loss of hemagglutination activity of Gal-8, which it possesses due to its bivalent carbohydrate binding capacity (32).. Here we aimed to determine if Gal-8 proteolysis by M^pro^ had an effect on its immune regulatory activity and cytokines secretion, given the fact that cytokines play essential roles in acute and chronic viral infections which can have beneficial effects during viral clearance.

### The cleavage of Gal-8 by M^pro^ affects its immunomodulatory activity

To assess if cleaved Gal-8 was able to demonstrate immune regulatory properties, FL-Gal-8 was incubated with M^pro^ overnight at 37 °C, and the protein sample was subjected to size-exclusion chromatography where Gal-8159-317 and Gal-810-158 products were separated from FL-Gal-8 and M^pro^. Following this human peripheral blood mononuclear cells (PBMCs) were cultured in the absence or presence of either FL-Gal-8 or Gal-8^159-317^ and Gal-8^10-158^ sample (1 µg/ml) overnight. We found that both samples of Gal-8 enhanced TNF-α and IL-6 production in the culture supernatants as measured by ELISA (Fig. 4A, B). However, the truncated Gal-8 displayed less pronounced immunomodulatory effects compared to the FL-Gal-8 (Fig. 4A and 4B).

**Figure 4.**
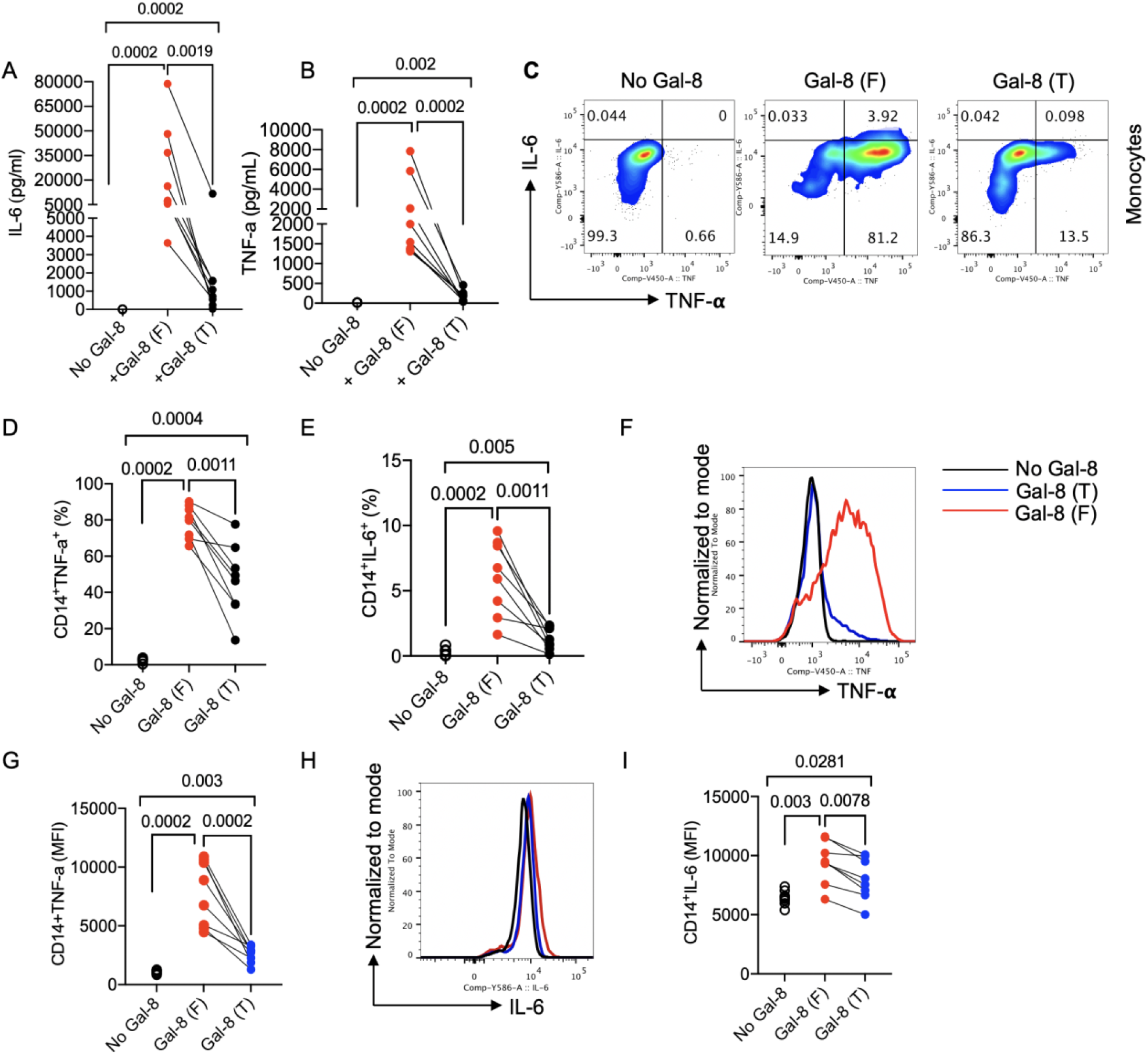
**(A)** Quantification of IL-6 and **(B)** TNF-α in culture supernatants following stimulation with the full length (F), or the truncated version of Gal-8 (T), consisting of Gal-8 ^10-158^ and Gal-8 ^159-317^, as measured by ELISA. **(C)** Representative flow cytometry plots, **(D)** cumulative data of percentages of TNF-α, and **(E)** percentages of IL-6 expressing cells among CD14+ monocytes. **(F)** Representative histogram plots, and **(G)** cumulative data of the mean fluorescence intensity (MFI) of TNF-α in CD14+ cells either unstimulated (No Gal-8) or treated with Gal-8 (F) and Gal-8 (T). **(H)** Representative histogram plots, and **(I)** cumulative data of the mean fluorescence intensity (MFI) of IL-6 in CD14+ cells either unstimulated (No Gal-8) or treated with Gal-8 (F) and Gal-8 (T). Each dot represents data from an individual human study subject.

To identify Gal-8 target cells, we subjected PBMCs to further analysis by flow cytometry. These studies revealed that 6 h treatment of PBMCs with both samples of Gal-8 significantly increased the percentages of TNF-α/IL-6 expressing cells among CD14+ monocytes in vitro (Figure 4C-E). We found that both full length and truncated forms of Gal-8 not only increased the frequency of TNF-α/IL-6 expressing monocytes but also elevated the intensity of TNF-α (Figure 4F, G) and IL-6 expression cells among these monocytes (Figure 4H, I). Nevertheless, the truncated version of Gal-8 demonstrated significantly less prominent stimulatory effects on monocytes than the full length (Fig. 4 C-I).

Moreover, we observed that Gal-8 exhibited a similar effect on B cells (CD19+ cells). As shown in Figure 5A-C, either the full or truncated version of Gal-8 significantly increased the proportion of TNF-α/IL-6 expressing cells among B cells. Similar to monocytes, the full length of Gal-8 had a more pronounced stimulatory effect on B cells than its truncated version (Figure 5A). However, neither FL nor truncated Gal-8 had any stimulatory effects on both CD4+ and CD8+ T cells in terms of cytokine production (data not shown).

**Figure 5.**
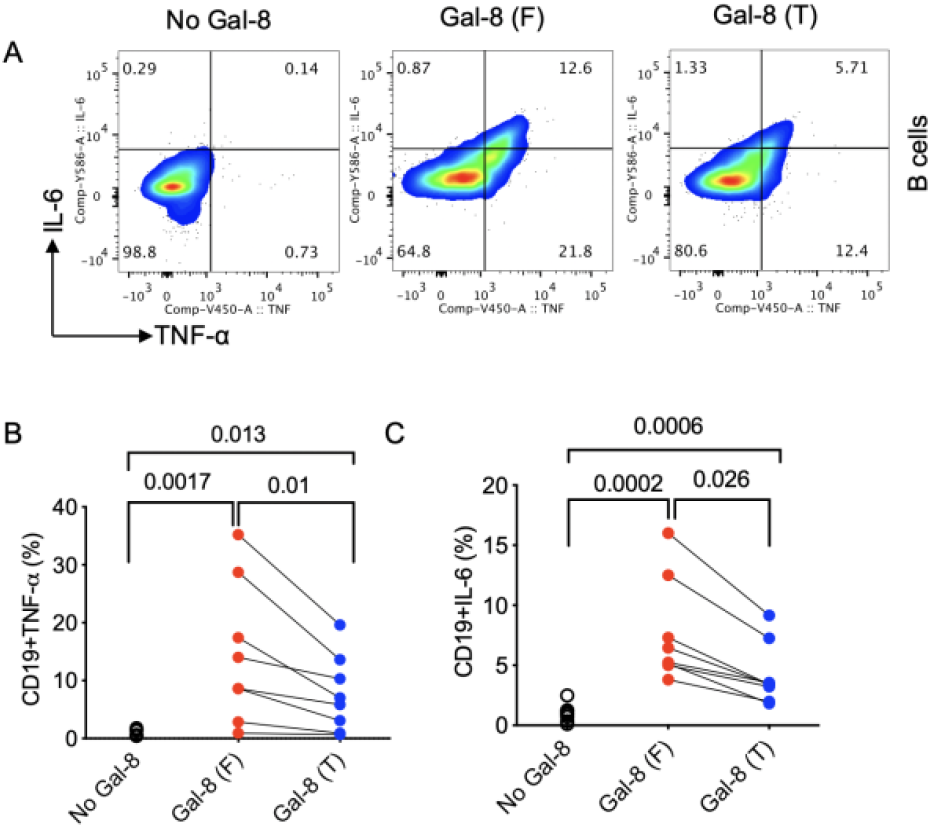
**(A)** Representative flow cytometry plots, **(B)** cumulative data of percentages of TNF-α, and **(C)** percentages of IL-6 expressing cells among CD19+ B cells.

Together this data suggests Gal-8 cleavage will have alterations in host immune response as a result of compromised/impaired cytokine stimulation. Importantly, we do note that increased Gal-8 cleavage was observed in M^pro^ mutations T21I, S46F, E47K and L50F and R279C found in the Delta SARS-CoV-2 VOC (Fig. 3E), which has been linked to IFN suppression and higher virulence (48). In clinical samples, a low production of IFN-γ was associated with more severe cases of COVID-19 (49), and accordingly the Delta VOC is able to produce high viral loads and increased transmission compared to other VOCs (50, 51).

## DISCUSSION

Understanding the pathophysiology of COVID-19 still remains a high priority, especially in the light of newly emerging VOCs. Every discovered variant presents a potential risk of possible clinical implications due to alterations and newly acquired properties in viral components leading to increased transmissibility and/or higher replication rate. This may result in resistance to existing medications or inhibitors under development. Therefore, tracing the mutations occurring in SARS-CoV-2 viral components is essential to ensure the efficacy of potential antivirals.

In this study we analysed 31 mutations of M^pro^ found in 5 VOCs of SARS-CoV-2. We identified hot spots that had an effect on protease functionality and structural integrity. The hot spots were located around or in close proximity to the active site or at the dimer interface – the region involved in allosteric regulation of M^pro^ activity. Interestingly, previous studies of M^pro^ from SARS-CoV and SARS-CoV-2 particularly demonstrated the importance of long-range interaction for protein structure and function where even a single point mutation might cause drastic effect for protease function (25, 52-54). However, since we selected mutations found in sequences of clinical isolates and taking into account that M^pro^ function is critical for virus life cycle, we did not expect to find mutations that would cause a significant negative effect on protease activity.

We employed crystallographic structural analysis together with high fidelity FRET activity assay as described previously by our group to assess the functional and structural changes in M^pro^ mutants (23, 55). The catalytic parameters for most substitutions were not significantly altered with the maximum level of change of 1.7-fold for catalytic efficiency and 2-fold for KM values. These findings were not surprising, given the recent study demonstrating that most of the residues, including those that form substrate binding pockets, were able to tolerate quite a bit of variability while maintaining functionality and structural integrity (56, 57). We observe the highest level of conformational plasticity around S2’ and S3’ binding pockets with N142 and S46 being the most flexible residues within the structures we solved. Multiple rotamers of N142 were also observed in previous studies (57, 58). N142 is known to have an important role in substrate binding and in binding of several tested inhibitors (24, 59). The flexible side chain of N142 extends over the S1 subsite, forming van der Waals interactions with Gln in P1 position (58). In several crystal structures of M^pro^ with inhibitors, N142 forms hydrogen bonds with water in the active site, stabilizing the bound inhibitor (24, 59). The plasticity of N142 may enhance the binding of Gln which contributes to the conserved feature of M^pro^ cleavage sites.

Also, we observed that the charge distribution in the active site changes quite dramatically even upon a single substitution especially around the S3’ and S2 binding pockets. This might explain why with most mutants we see the changes in binding affinities and not the kcat values. The active site of M^pro^ is unique because it has a natural ability of being malleable enough to accommodate 11 similar yet slightly different cleavage sequences of the viral polypeptide. Several mutations we explored (L50F and E47K/N) were located in close proximity to important elements of the active site and significantly changed the surface charge and orientation of surrounding residues, which were involved in S2, S3’ and S4 pockets. The S2 subsite has a preference for Lue but also the plasticity to accommodate Phe as P2 in the 11 endogenous cleavage sequences (60). Crystal structures of apo M^pro^ reveal that S2 subsite adopts a more open conformation in the empty active site. Upon the binding of Leu in P2 position, the side chains of M49 and Q189 are redirected, inducing conformational changes in S2 subsite. L50F and E47N are the mutants that cause the most profound changes in S2 (Figure 2B, C). L50 is located at the surface of the active site, which might contribute to the entrance of substrate binding groove with an open conformation (57). The substitution of leucine with a bulkier aromatic phenylalanine resulted in a wider open S2 pocket compared to the wild type. Meanwhile M49, M165 and Q189 are positioned to form additional hydrogen bonds to narrow S2 in E47N active site. The conformational changes of S2 in structures of apo M^pro^ mutants might lead to sufficient or insufficient binding of substrates indicated by the 3.5-fold difference in KM values of L50F and E47N. Despite this, the changes in catalytic parameters were not significant compared to the wild type (Figure 1C, D).

Another group of mutations of interest was located at the dimer interface (Figure 1A). M^pro^ has an allosterically regulated correlation between dimerization and catalysis, demonstrated by the effect of mutating residues involved in dimerization on activity (52, 61, 62). However, the apo-structures of the corresponding mutants did not reveal any significant structural changes. Activity assays also did not result in functional consequences.

To further assess if the structural plasticity of the active site of M^pro^ might be affected by mutations, we performed substrate specificity studies. Since the structures of several mutants revealed altered orientations of side chains of residues involved in substrate binding pockets and reorganization of hydrogen bond networking around the active site, the M^pro^ ability to recognize certain amino acids could be affected as well. HyCoSuL approach demonstrated that several mutants indeed had an altered specificity (Figure 2G). The most interesting changes we observed were C-terminus mutants A260V and S284G that became more permissive at the P2 position. The most preferred amino acid in this position is Leu for the wild type, but other hydrophobic residues can be accommodated for the mutants, however with lower activity. The A260V mutant displayed activity with almost all natural amino acids at P2. Looking at the structures of S284G and A260V mutants, we noticed that the S2 binding pocket and the area around it have lost negative charge in both cases (Figure S2D, L). This might explain the difference in specificity in that region. Dimer interface consisting of the N-finger and C-terminal helix plays a major role in allosteric communication in the M^pro^ dimer (63). Dimerization or dimer packing can influence the stabilization of both the active site and the protein; therefore, apo-structures may lack distinctive conformational changes to explore the substrate specificity of C-terminal mutants. Further study could assess the distal regions by solving and comparing the ligand-bound crystal structures.

Viral proteins are evolutionary evolved to be multifunctional and the promiscuous nature of substrate specificity of M^pro^ can be explained by its pleotropic role. It was logical to assume that the viral protease is able to cleave not only viral substrates but also host proteins, adding to its complex role in infection. Recently, N-terminomics studies identified more than 100 M^pro^ substrates in human lung and kidney cells (32). Gal-8 – a host defence protein and one of the key regulators of immune response – was shown to be one of the host substrates. Gal-8 is responsible for secretion of cytokines and chemokines and involved in the development of cytokine storm, which was also reported for Gal-9 (64) – a dangerous condition associated with more severe COVID-19 outcomes. In this study, we confirmed the cleavage of Gal-8 by M^pro^ and identified at least two additional cleavage sites by mass spectrometry – the one that was identified previously (Q158 in a linker region) and two additional sites at the termini, Q^9^ and Q^248^. The cleavage of Gal-8 by M^pro^ therefore follows more complex enzyme kinetics with several cleavage sites involved, some substrate products being intermediate (34 kDa and 21 kDa bands) and several products being stable after 24 hours of cleavage, such as Gal-810-158 and Gal-8^159-317^. Interestingly, SARS-CoV M^pro^ demonstrated a similar pattern and rate for Gal-8 substrate cleavage. The stable cleavage product Gal-8^159-317^ was monitored while comparing the activity among variants. Even though the difference in activity between M^pro^ variants towards Gal-8 was quite noticeable it remains to be determined how M^pro^ function relates to viral fitness and pathogenicity, and it is possible that in order to see the significant consequences for the host immune system the functionality of a variant’s M^pro^ needs to be altered by a large amount. However, the known outcome of Gal-8 cleavage is the loss of ability to recruit the autophagy adaptor NDP52 to damaged endosomes, which allows SARS-CoV-2 to escape antiviral xenophagy. It was confirmed by immunoprecipitation that NDP52 binds the C-terminal domain of Gal-8 after its cleavage, but if Gal-8 is cleaved further at the newly identified Q246 site Gal-8 interaction with NDP52 might be also prevented.

In addition to the direct interaction of cytosolic Gal-8 with pathogen proteins, secreted Gal-8 also plays a role in the regulation of adaptive and innate immune responses by targeting cytokine receptors and inducing TNF signaling and cytokines and chemokines expression. Given the important role of Gal-8 in pathogenicity we sought to assess if M^pro^ cleavage affected the immunomodulatory properties of Gal-8. Interestingly, the cleavage products Gal-8^10-158^ and Gal-8^159-317^ still were able to enhance TNF-α/IL-6 expression in both PBMC culture and in CD14+ monocytes and B cells in vitro but with a less profound effect compared to the full-length of Gal-8, contributing to decreased level of host immunocompetence. Thus, the single event of M^pro^-mediated cleavage of Gal-8 influences multiple ways for the virus to evade host defence pathways. First, intracellularly it defeats antiviral mechanism and allows SARS-CoV-2 to escape xenophagy, as it was demonstrated before (32) and secondly once secreted the truncated form of Gal-8 causes a decreased immune response effect against viral infection. Further in vivo studies need to be performed to investigate the physiological effects of Gal-8 cleavage by different M^pro^ mutants and VOCs.

Lastly, emerging new variants cause a valid concern that Covid-19 might develop a resistance to known antivirals through mutation of vital amino acids residues in the active site important for drug binding. Importantly, despite some structural changes in substrate binding pockets for several mutants as well as alterations in KM values for viral substrate we still observed maintained potency of nirmatrelvir and our derivative inhibitor against the variants of M^pro^, suggesting M^pro^ remains an excellent antiviral target as the virus evolves

As the COVID-19 pandemic continues to pose a global health threat with the increased ability of variants to spread and escape immune responses, SARS-CoV-2 antiviral drugs are necessary to combat the pandemic and to prevent future outbreaks. Potential drug resistance needs to be considered for the inhibitors targeting M^pro^ mutants with altered catalytic properties on viral and host substrates, so it is crucial to investigate the potency of existing antivirals. Future variants require close monitoring for possible drug resistance which needs to be considered for the development of next-generation M^pro^ inhibitors.

FIGURES (Word Style “VA_Figure_Caption”). Each figure must have a caption that includes the figure number and a brief description, preferably one or two sentences. The caption should follow the format “Figure 1. Figure caption.” All figures must be mentioned in the text consecutively and numbered with Arabic numerals. The caption should be understandable without reference to the text. Whenever possible, place the key to symbols in the artwork, not in the caption. To insert the figure into the template, be sure it is already sized appropriately and paste before the figure caption. For formatting double-column figures, see the instructions at the end of the template. Do NOT modify the amount of space before and after the caption as this allows for the rules, space above and below the rules, and space above and below the figure to be inserted upon editing.

## DATA AND MATERIALS AVAILABILITY

All data are available in the main text or the supplementary materials. Inhibitors are available with materials transfer agreements (MTAs). Structural data has been deposited in the www.rscb.org database with accession numbers: 8DJJ.PDB M^pro^-A7T; 8EJ9.PDB: M^pro^-E47N; 8EJ7.PDB: M^pro^-E47K; 8DKZ.PDB: M^pro^-L50F; 8DKL.PDB: M^pro^-L89F; 8DKJ.PDB: M^pro^-K90R; 8DI3.PDB: M^pro^-P132H; 8DK8.PDB: M^pro^-T190I.

## ASSOCIATED CONTENT

### Supporting Information

Supporting Information includes Methods and Supplemental figures. This material is available free of charge via the Internet at http://pubs.acs.org.“

## AUTHOR INFORMATION

### Author Contributions

Conceptualization: EA, SAC, JL, MJL

Methodology: SAC, JL, EA, MBK, JI, EM, WR, MZ, SS, ZT, BB, TL

Investigation: SAC, JL, MBK, JI, EM, MZ, WR, SS, ZT, BB, TL

Visualization: EA, SAC, JL

Supervision JAN, OJ, SE, JN, MD, HSY, JCV, MJL

Writing—EA, SAC, JL

Writing—review & editing: SAC, EA, JL, MBK, WR, SS, JI, EM, ZT, BB

### Funding Sources

Canadian Institutes of Health Project grant PJT 180390 Canadian Institutes of Health Research Rapid Research (VR3-172655).

Natural Sciences and Engineering Research Council of Canada (549297-2019)

Alberta Innovates

Alberta Jobs, Economy and Innovation Infrastructure Grant - Government of Alberta Foundation for Polish Science

## ACKNOWLEDGMENT

We thank the staff at SSRL beamline 12-1, in particular Dr. Silvia Russi. Use of the Stanford Synchrotron Radiation Lightsource, SLAC National Accelerator Laboratory, is supported by the U.S. Department of Energy, Office of Science, Office of Basic Energy Sciences under Contract No. DE-AC02-76SF00515. The SSRL Structural Molecular Biology Program is supported by the DOE Office of Biological and Environmental Research, and by the National Institutes of Health, National Institute of General Medical Sciences (P30GM133894).

## ABBREVIATIONS

M^pro^: Main protease
Gal-8: Galectin-8

---

SYNOPSIS TOC (Word Style “SN_Synopsis_TOC”). If you are submitting your paper to a journal that requires a synopsis graphic and/or synopsis paragraph, see the Instructions for Authors on the journal’s homepage for a description of what needs to be provided and for the size requirements of the artwork.

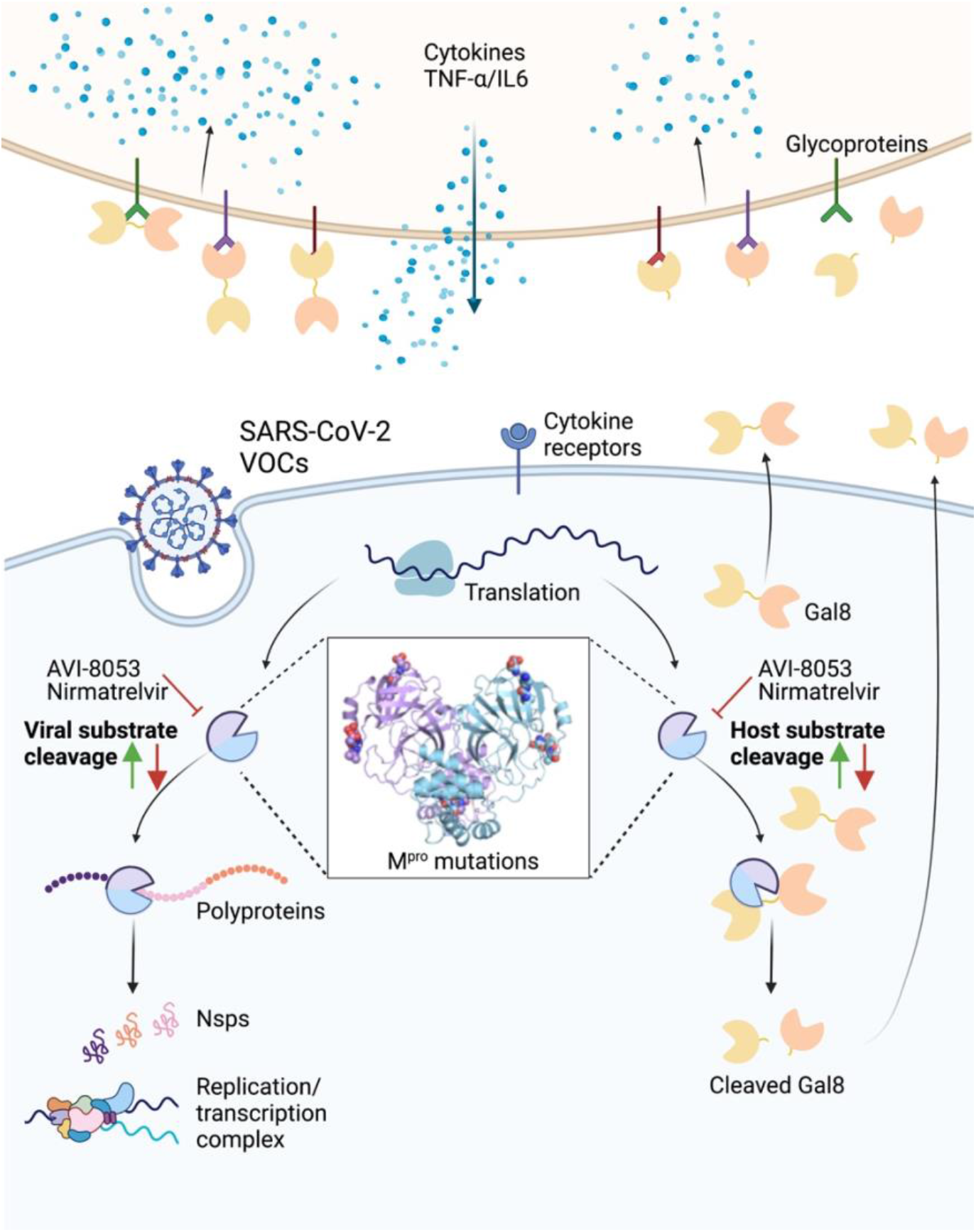

