## Supplementary material for "SARS-CoV-2 M^pro^ protease variants of concern display altered viral and host target processing but retain potency towards antivirals": Supplementary_Materials BIORxiv.pdf

**This PDF file includes:**

Figures. S1 to S9
Tables S1 to S2

**Table of Contents:**

1. **Methods** **S2-S7**
2. **References** **S8**
3. **Supplemental figures** **S9-S26**

**Methods**

**Mutation identification and prevalence calculations.**

Only "high coverage" sequences were selected for analysis. The prevalence was calculated as a percentage of a certain mutation compared to the amount of all mutations in *nsp5* gene in a particular variant.

**Mutagenesis and protein purification.**

**SARS-CoV-2 $M^{pro}$.** $M^{pro}$ SARS-CoV-2 gene with N-terminal His-SUMO tag in pET SUMO expression vector (Invitrogen) was used for protein production. The mutations were introduced into protein gene using Site-Directed mutagenesis kit (Agilent, Canada). Purification of SARS-CoV-2 $M^{pro}$ wild type and variants were performed as described earlier (*1*).

**Gal-8.** Gal-8-6X His gene in pET vector was transformed into Rosetta (DE3) *E.coli* cells and the protein was expressed for 18 hours at 18 °C. The harvested cells were resuspended in 50 mM HEPES, pH 7.0, 300 mM NaCl, 5 mM Imidazole buffer, lysed by sonication and the protein was purified using.

**Gal-9.** Gal-9-6X His gene in pBad vector was transformed into TOP-10 *E. coli* cells and the protein was expressed for18 hours at 18 °C. The harvested cells were resuspended in 50 mM Tris-HCl, pH 7.5, 300 mM NaCl, 5% glycerol, 5mM Imidazole buffer and lysed by sonication followed by d centrifugation step at 17,000 g for 30 min and Ni-NTA affinity chromatography. The final protein sample was concentrated using Amicon Ultra-15 (MWCO 10 kDa) and flash frozen.

**HyCoSuL screening**

Substrate specificity profile of SARS-CoV-2 $M^{pro}$ wild-type and the mutants was determined in the same manner as described previously (*2*). Library screening was performed using a spectrofluorometer (Molecular Devices Spectramax Gemini XPS) on 96-well plates. Enzymes were diluted in assay buffer (20 mM Bis-Tris, 1 mM DTT, pH 7.4) and added to plate wells. Assay conditions were 1 µL of substrate in DMSO and 99 µL of enzyme. Substrate hydrolysis was measured for 60 min at the appropriate wavelength ($\lambda_{ex}$ = 355 nm, $\lambda_{em}$ = 460 nm).

**FRET-based SARS-CoV-2 $M^{pro}$ activity assay.**

FRET-based cleavage assays with synthesized fluorescent substrate of SARS-CoV-2 $M^{pro}$ (Abz-SVTLQ↓SG-Tyr($NO_2$)-R) were conducted as described previously (*3*).

**SARS-CoV-2 $M^{pro}$ activity assay with Gal-8.**

Gel-based end point activity assay using recombinant Gal-8 was conducted using 14 µM of substrate and 2.5 µM of SARS-CoV-2 $M^{pro}$ WT or variants in activity buffer (20 mM Bis–Tris, pH 7.8, 1 mM DTT). The mixture was incubated at 37 °C, 30 µL aliquots were taken out at specific time points, the reaction was stopped by adding 5 µM of GC376 inhibitor and then SDS-sample buffer. Protein samples were analysed by SDS-PAGE and digitization was carried out with ImageQuant LAS 4000 (GE Healthcare, USA).

**Differential scanning fluorimetry.**

DFS was performed using 6 µM of protease with a final concentration of 5X for SyproOrange dye (Thermo Fisher Scientific, USA) in 50 mM Tris-HCl, pH 8.0, 150 mM NaCl. All samples were run in duplicate. The thermal scan was conducted from 25 to 95 °C, at 0.5 °C/min (ViiA 7 Real-Time PCR System, ThermoFisher). The melting point ($T_m$) was calculated by fitting the raw fluorescence data over the temperature using the Boltzmann equation in GraphPad Prism program (Prism 9, GraphPad Software, USA).

**Human subject's ethics statement.**

This study was approved by the Research Ethics Board at the University of Alberta (protocol # Pro000046064). A written informed consent form was obtained from the study participants.

**Sample processing.**

Peripheral blood mononuclear cells (PBMCs) were isolated from the blood using Ficoll-Paque gradients. Cell cultures were performed in RPMI 1640 (Sigma-Aldrich) supplemented with 10% FBS (Sigma-Aldrich) and 1% penicillin/streptomycin (Sigma-Aldrich).

**Cell culture and Flow cytometry.**

Human cell antigens and cytokines fluorophore-labelled antibodies were purchased from BD Biosciences and Thermo Fisher Scientific and used for anti-CD3 (SK7), anti-CD4 (RPA-T4), anti-

CD8 (RPA-T8), anti-CD14 (61D3), anti-CD19 (HIB19), anti-TNF-α (MAB11), and anti-IL-6 (MQ2-13A5). Live dead staining was used to exclude dead cells. Surface and intracytoplasmic cytokine staining (ICS) was performed according to our previous reports (*4-6*). For ICS, PMBCs were cultured and stimulated with LPS (1 µg/mL), Gal-8 full length or truncated (1 µl/ml) for 6 hr in the presence of Brefeldin A (1 µl/ml). Cells were fixed and permeabilized and acquired on an LSRFortessa-SORP and analyzed using the FlowJo software (version 10).

**ELISA**

PBMCs were cultured in the absence or presence of Gal-8 full length or truncated (1 µl/ml) for overnight. The concentration of TNF-α in the culture supernatants was measured using an ELISA kit (R&D Systems).

**Statistical analysis**

The non-parametric tests such as the Mann-Whitney U-test or Kruskal–Wallis one-way analysis of variance was used. The P-values are shown in the graphs and measures are expressed as mean ± SEM and P-value < 0.05 was considered to be statistically significant.

**Crystallography**

SARS-CoV-2 $M^{pro}$ wild-type and variants were dialyzed against 5 mM Tris-HCl pH 8.0, 10 mM NaCl buffer overnight at 4 °C. Protein was concentrated to 8 mg/ml. SARS-CoV-2 $M^{pro}$, wild-type and variants, were subjected to the JCSG plus and PACT crystallization screen (Molecular Dimensions) with hits identified under several conditions. The best crystals for SARS-CoV-2 $M^{pro}$ A7T and K90R were observed with sitting drop at room temperature at a ratio of 1:1 with mother liquor 0.2 M lithium chloride, 0.1 M HEPES pH 7.0 and 20% w/v PEG6000. The best crystals for SARS-CoV-2 $M^{pro}$ E47N were observed with sitting drop at room temperature at a ratio of 1:1 with mother liquor 0.2 M sodium fluoride, 20% w/v PEG3350. The best crystals for SARS-CoV-2 $M^{pro}$ E47K was observed using sitting drop at room temperature at a ratio of 1:1 with mother liquor 0.2 sodium thiocyanate, 0.1 M Bis-Tris propane pH 7.5, 20% w/v PEG3350. The best crystals for SARS-CoV-2 $M^{pro}$ L50F was observed using sitting drop at room temperature at a ratio of 1:1 with mother liquor 0.1 M MMI pH 9.0, 25% w/v PEG1500. The best crystals for SARS-CoV-2 $M^{pro}$ L89F were observed using sitting drop at room temperature at a ratio of 1:1

with mother liquor 0.2 M sodium formate, 0.1 M Bis-Tris Propane pH 6.5, 20% w/v PEG3350. The best crystals for SARS-CoV-2 M$^{pro}$ P132H were observed using sitting drop at room temperature at a ratio of 1:1 with mother liquor 0.2 ammonium phosphate monobasic, 0.1 M Tris pH 8.5, 50% v/v MPD. The best crystals for SARS-CoV-2 M$^{pro}$ T190I, A260V, G283S and S384G were observed using sitting drop at room temperature at a ratio of 1:1 with mother liquor 0.2 M sodium chloride, 0.1 M HEPES pH 7.0, 20% w/v PEG6000. Prior to freezing, crystals were incubated with 22% glycerol as a cryoprotectant. Data collection took place at Stanford Synchrotron Radiation Lightsource (SSRL) (Menlo Park, CA) beamline 12-2 with Blu-Ice using the Web-Ice interface and at Canadian Light Source, Inc. (CLSI) beamline 08B1-1 using MxDC.

**Diffraction data collection, phase determination, model building, and refinement**

Diffraction data sets for SARS-CoV-2 M$^{pro}$ A7T, L50F and T190I were collected at 100 K in a cold nitrogen stream using CLSI beamline 08B1-1 coupled to PILATUS 6M. Diffraction data sets for SARS-CoV-2 M$^{pro}$ E47N, E47K, L89F, K90R, T190I, A260V, G283S and S284G were collected at 100K in a cold nitrogen stream using SSRL beamline 12-2 at a wavelength of 0.979460, equipped with an Eiger 16 M Pixel Array detector. Multiple data sets were collected per variant with the best selected based on data statistics. X-ray detector Software (XDS) and Scala were used for processing of the data sets. All structures were determined by molecular replacement, with the crystal structure of the free enzyme of the SARS-CoV-2 M$^{pro}$ (PDB entry 6WTM) as a search model, using the Phaser program from Phenix, version v1.19.2-4158. Refinement of all eleven structures were performed with phenix.refine in Phenix software. Statistics of diffraction, data processing and model refinement are given in Supplementary Table 3. The model was inspected with Ramachandran plots and final models displayed using PyMOL molecular graphics software (Version 2.5.2 Schrödinger, LLC.)

**In-gel Trypsin Digestion**

SARS-CoV-2 M$^{pro}$ and Gal-8 were separated after the cleavage assays by 14% SDS-PAGE. The gels were fixed for 20 minutes (50% ethanol, 2% phosphoric acid), washed twice for 20 minutes each ($ddH_2O$) and stained overnight with blue-sliver coomassie stain (20% ethanol, 10% phosphoric acid, 750 mM ammonium sulphate, 0.12% Coomassie Blue G-250) and washed twice for 10 minutes each ($ddH_2O$). Each lane was separated into 2 fractions and cut into 1 mm pieces for analysis. The gel bands were transferred to a round bottom 96-well plate and to each well 200 µl of destaining solution (50 mM ammonium bicarbonate, 50% acetonitrile) was added. The plate was incubated at 37 °C for 10 minutes, the solution was removed, and the distaining was repeated 3 times and then the solution was replaced with acetonitrile. The plate was incubated again at 37 °C for 10 minutes. The dehydration was repeated until the gel bands became white (2 times) and the sample were dried at 37 °C for 10 minutes. The gel bands were rehydrated with 200 µl of reducing solution (100 mM ammonium bicarbonate, 11.4 mM beta-mercaptoethanol) and incubated at 37 °C for 30 minutes. The reducing solution was removed, 200 µl of alkylating solution (100 mM ammonium bicarbonate, 10 mg/mL iodoacetamide) was added, the gel bands were incubated at 37 °C for 30 minutes and washed with 200 µl of 100 mM ammonium bicarbonate at 37 °C for 10 minutes twice, dehydrated in 200 µl acetonitrile at 37 °C for 10 minutes and dried at 37 °C for 15 minutes. The gel bands were trypsinized (100 µl of 100 mM ammonium bicarbonate and 6 µg/ml Sequence Grade Modified Trypsin, Promega Inc.) overnight. The solutions containing tryptic peptides were transferred to a round bottom 96-well plate. Tryptic peptides were further extracted from the gel bands with extraction solution (2% acetonitrile, 1% formic acid) and incubated at 37 °C for 1 hour. A final extraction was conducted using 50% acetonitrile and 0.5% formic acid and incubated at 37 °C for 1 hour. All solutions containing tryptic peptides were transferred to a round bottom 96-well plate and freeze-dried under vacuum overnight. The samples were resuspended (1 fraction per lane) in 0.1% formic acid before analysis by liquid chromatography tandem mass spectrometry (LC-MS/MS).

**Mass spectrometry analysis**

Samples were analyzed using a nanoflow-HPLC (Thermo Scientific EASY-nLC 1200 System) coupled to an Orbitrap Fusion Lumos Tribrid Mass Spectrometer (Thermo Fisher Scientific). The peptide mixture underwent reverse phase separation by an analytical column (2 µm, 100 Å, 50 µm × 15 cm, PepMap RSLC C18: Thermo Fisher Scientific). Peptides were eluted over a 45 minute (whole assay) or a 30 minute (individual protein fragment bands) linear gradient from 0 to 36.8% acetonitrile in 0.1% formic acid. Data analysis was conducted using ProteinProspector (v5.22.1) against a concatenated database of the Homo sapien proteome (SwissProt.2017.11.01. random.concat). The search parameter included digestion by TrypsinPro with non-specificity at the N-termini, a maximum of 3 missed trypsin cleavages, a precursor change range of +2 +3 +4, a precursor mass tolerance of 15 ppm, a fragment mass tolerance of 0.8 Da, carbamidomethylation of Cys (constant modification), and oxidation of Met and deamidation of Asn and Gln (variable modifications). A decoy database search was conducted to evaluate the false-positive rates. The target false discovery rate was set at 2% at the protein level and 1% at the peptide level.

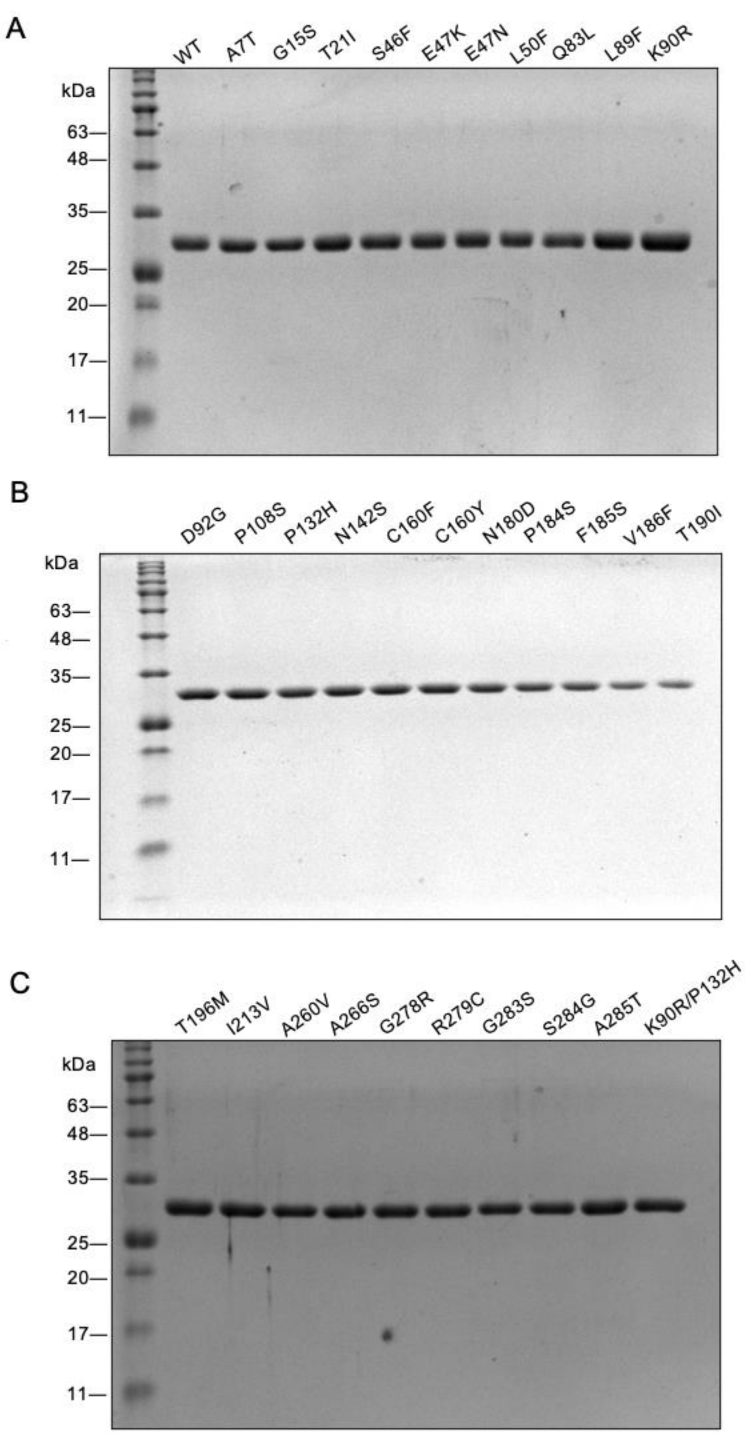


**Figure S1.** SDS-PAGE of recombinant SARS-CoV-2 $M^{pro}$ wild-type and the mutants. (A-C). The wild type and the VOC mutants of $M^{pro}$ were purified using Ni-NTA resin, followed by SUMO tag removal and Size Exclusion Chromatography on Superdex75 (10/300) column.

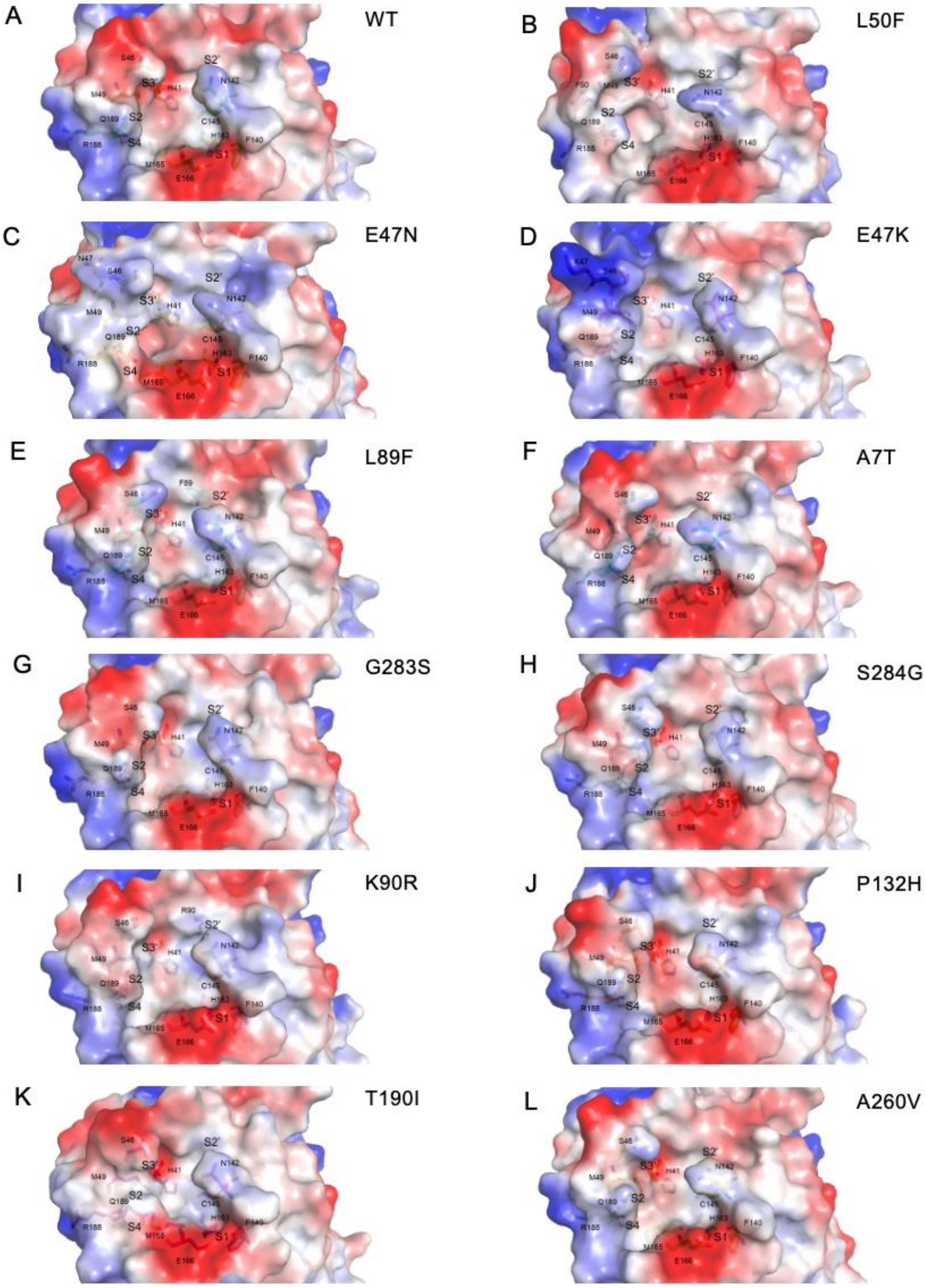


**Figure S2.** Molecular surface representations showing the electrostatic surface potentials of $M^{pro}$ wild type (A) and the mutants (B-L). All residues in the active site pocket are labelled.

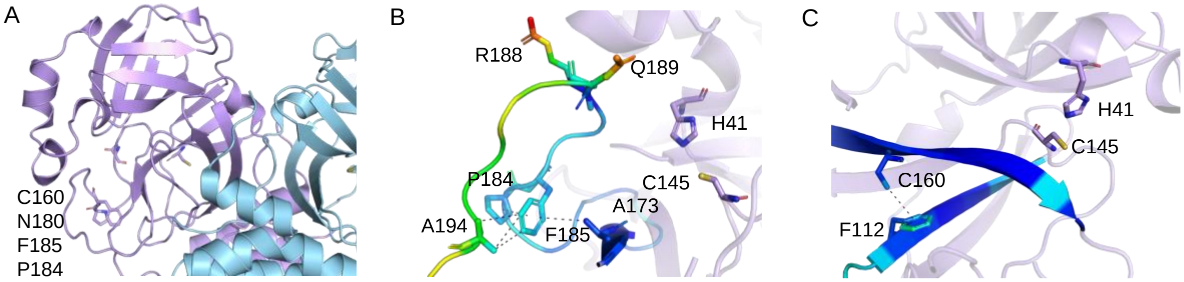


**Figure S3.** The distribution of mutations with most changes in thermal stability in SARS-CoV-2 M$^{pro}$ molecule (PDB 6WTM). (A) The location of mutations in M$^{pro}$ molecule, which demonstrated the most drastic changes in thermal stability assessed by DSF. (B) and (C) B-factor differences are displayed for the structural elements in a region of M$^{pro}$ molecule where destabilizing mutations were found as a blue–yellow–green color ramp with blue indicating the smallest value (stabilization) and green indicating the largest value (destabilization).

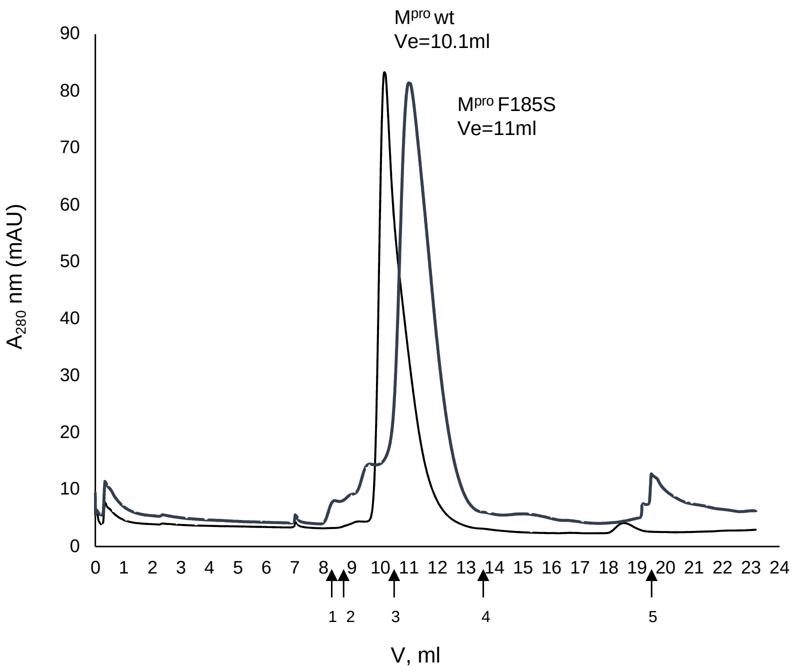


**Figure S4.** Size exclusion chromatography of M$^{pro}$ wt (black) and M$^{pro}$ F185S (grey) performed on Superdex 75 Increase (10/300) column (Cytiva). Standards: thyroglobin, 8.38 ml, 670 kDa (1. ); γ-globulin, 8.78 ml, 158 kDa (2); ovalbumin, 10.67 ml, 44 kDa (3); myoglobin, 13.46 ml, 17 kDa (4); vitamin B12, 19.34 ml, 1.35 kDa.

Ac-Mix-P3-Mix-Gln-ACC

| Variants | L-Ala | L-Arg | L-Asn | L-Asp | L-Glu | L-Gln | Gly | L-His | L-Ile | L-Leu | L-Lys | L-Phe | L-Pro | L-Ser | L-Thr | L-Trp | L-Tyr | L-Val |
|---|---|---|---|---|---|---|---|---|---|---|---|---|---|---|---|---|---|---|
| WT | 5 | 22 | 0 | 0 | 0 | 15 | 0 | 7 | 14 | 15 | 33 | 0 | 0 | 0 | 18 | 11 | 8 | 23 |
| E47K | 3 | 15 | 0 | 0 | 0 | 0 | 0 | 0 | 15 | 11 | 22 | 0 | 0 | 0 | 21 | 10 | 6 | 26 |
| E47N | 0 | 21 | 0 | 0 | 0 | 13 | 0 | 8 | 19 | 14 | 38 | 0 | 0 | 0 | 18 | 12 | 7 | 31 |
| L50F | 0 | 0 | 0 | 0 | 0 | 8 | 0 | 0 | 17 | 13 | 22 | 0 | 0 | 0 | 11 | 18 | 0 | 32 |
| L89F | 0 | 17 | 0 | 0 | 0 | 0 | 0 | 0 | 17 | 13 | 39 | 0 | 0 | 0 | 17 | 12 | 6 | 27 |
| K90R | 0 | 29 | 0 | 0 | 0 | 22 | 0 | 0 | 19 | 23 | 42 | 0 | 0 | 0 | 25 | 19 | 16 | 38 |
| P108S | 0 | 0 | 0 | 0 | 0 | 0 | 0 | 0 | 18 | 12 | 31 | 0 | 0 | 0 | 15 | 10 | 3 | 24 |
| P132H | 0 | 19 | 0 | 0 | 0 | 0 | 0 | 0 | 16 | 12 | 35 | 0 | 0 | 0 | 17 | 13 | 9 | 24 |
| A260V | 0 | 9 | 0 | 0 | 0 | 8 | 0 | 0 | 16 | 11 | 21 | 0 | 0 | 0 | 19 | 10 | 0 | 22 |
| G283S | 14 | 13 | 0 | 0 | 0 | 0 | 0 | 0 | 14 | 8 | 29 | 0 | 0 | 0 | 14 | 15 | 0 | 21 |
| S284G | 0 | 15 | 0 | 0 | 0 | 7 | 0 | 0 | 10 | 12 | 27 | 0 | 0 | 0 | 17 | 10 | 8 | 21 |

Ac-P4-Mix-Mix-Gln-ACC

| Variant | L-Ala | L-Arg | L-Asn | L-Asp | L-Glu | L-Gln | Gly | L-His | L-Ile | L-Leu | L-Lys | L-Phe | L-Pro | L-Ser | L-Thr | L-Trp | L-Tyr | L-Val |
|---|---|---|---|---|---|---|---|---|---|---|---|---|---|---|---|---|---|---|
| WT | 80 | 0 | 0 | 0 | 0 | 0 | 0 | 0 | 11 | 0 | 0 | 0 | 31 | 29 | 55 | 27 | 0 | 85 |
| E47K | 93 | 0 | 0 | 0 | 0 | 0 | 0 | 0 | 0 | 0 | 0 | 0 | 0 | 21 | 35 | 14 | 0 | 82 |
| E47N | 73 | 0 | 0 | 0 | 0 | 0 | 0 | 0 | 0 | 0 | 0 | 0 | 13 | 23 | 40 | 15 | 0 | 69 |
| L50F | 52 | 0 | 0 | 0 | 0 | 0 | 0 | 0 | 0 | 0 | 0 | 0 | 0 | 0 | 16 | 0 | 0 | 53 |
| L89F | 65 | 0 | 0 | 0 | 0 | 0 | 0 | 0 | 0 | 0 | 0 | 0 | 27 | 31 | 42 | 25 | 0 | 69 |
| K90R | 77 | 0 | 0 | 0 | 0 | 0 | 0 | 0 | 0 | 0 | 0 | 0 | 0 | 0 | 0 | 0 | 0 | 84 |
| P108S | 65 | 0 | 0 | 0 | 0 | 0 | 0 | 0 | 0 | 0 | 0 | 0 | 0 | 0 | 30 | 16 | 0 | 61 |
| P132H | 89 | 0 | 0 | 0 | 0 | 0 | 0 | 0 | 0 | 0 | 0 | 0 | 15 | 23 | 38 | 13 | 0 | 92 |
| A260V | 55 | 0 | 0 | 0 | 0 | 0 | 0 | 0 | 0 | 0 | 0 | 0 | 0 | 0 | 21 | 0 | 0 | 55 |
| G283S | 46 | 0 | 0 | 0 | 0 | 0 | 0 | 0 | 0 | 0 | 0 | 0 | 0 | 22 | 33 | 17 | 0 | 53 |
| S284G | 59 | 0 | 0 | 0 | 0 | 0 | 0 | 0 | 0 | 0 | 0 | 0 | 18 | 13 | 35 | 15 | 0 | 58 |

Relative activity, %

0 20 100

**Figure S5.** Substrate specificity profile of SARS-CoV-2 M$^{pro}$ presented as heat maps using substrate library with natural amino acids assessing the P3 and P4 positions with Ac-Mix-P3-Mix-Gln-ACC and Ac-P4-Mix-Mix-Gln-ACC.

Ac-Mix-Mix-P2-Gln-ACC

| Variant | D-Ala | D-Arg | D-Asn | D-Asp | D-Gln | D-Glu | D-His | D-Leu | D-Lys | D-Phe | D-Pro | D-Ser | D-Phg | D-Thr | D-Trp | D-Tyr | D-Val | D-hPhe | B-Ala | L-Aze |
|---|---|---|---|---|---|---|---|---|---|---|---|---|---|---|---|---|---|---|---|---|
| WT | 0 | 8 | 0 | 0 | 8 | 0 | 0 | 0 | 0 | 0 | 0 | 0 | 0 | 0 | 6 | 0 | 0 | 7 | 5 | 0 |
| E47K | 0 | 6 | 0 | 0 | 0 | 0 | 0 | 0 | 0 | 0 | 0 | 0 | 0 | 0 | 7 | 0 | 0 | 0 | 10 | 0 |
| E47N | 0 | 0 | 0 | 0 | 0 | 0 | 4 | 5 | 5 | 0 | 6 | 6 | 0 | 0 | 6 | 0 | 0 | 0 | 9 | 0 |
| L50F | 0 | 0 | 0 | 0 | 0 | 0 | 0 | 5 | 0 | 4 | 0 | 0 | 0 | 0 | 8 | 0 | 0 | 6 | 9 | 0 |
| L89F | 0 | 0 | 0 | 0 | 0 | 0 | 0 | 4 | 0 | 0 | 0 | 0 | 0 | 0 | 5 | 0 | 0 | 0 | 8 | 0 |
| K90R | 0 | 0 | 0 | 0 | 0 | 0 | 6 | 8 | 8 | 7 | 9 | 7 | 6 | 0 | 8 | 6 | 6 | 6 | 13 | 0 |
| P108S | 0 | 8 | 0 | 0 | 8 | 0 | 6 | 7 | 0 | 0 | 0 | 0 | 0 | 0 | 8 | 0 | 0 | 0 | 14 | 0 |
| P132H | 0 | 0 | 0 | 0 | 0 | 0 | 0 | 0 | 0 | 0 | 0 | 0 | 0 | 0 | 11 | 0 | 0 | 11 | 16 | 0 |
| A260V | 7 | 11 | 9 | 4 | 10 | 6 | 8 | 9 | 8 | 6 | 10 | 8 | 7 | 5 | 10 | 7 | 6 | 9 | 16 | 6 |
| G283S | 4 | 6 | 5 | 0 | 4 | 11 | 9 | 9 | 4 | 0 | 6 | 8 | 8 | 0 | 9 | 4 | 0 | 3 | 11 | 0 |
| S284G | 0 | 8 | 6 | 4 | 7 | 0 | 5 | 6 | 5 | 6 | 9 | 5 | 7 | 4 | 9 | 0 | 6 | 9 | 13 | 5 |

| Variant | L-Hyp | L-Hyp(Bzl) | L-Thz | L-Oic | L-Idc | L-Pip | L-Tic | AC5C | dhAbu | dhLeu | L-Dap | L-Dab | L-Dab(Z) | L-Cit | L-hCit | L-Orn | L-Lys(TFA) | L-Lys(Ac) | L-Lys(2-ClZ) | L-Agp |
|---|---|---|---|---|---|---|---|---|---|---|---|---|---|---|---|---|---|---|---|---|
| WT | 0 | 6 | 5 | 0 | 5 | 6 | 6 | 0 | 2 | 3 | 0 | 0 | 0 | 2 | 6 | 0 | 1 | 0 | 0 | 3 |
| E47K | 0 | 8 | 0 | 7 | 11 | 0 | 6 | 0 | 0 | 0 | 0 | 6 | 0 | 0 | 0 | 0 | 0 | 0 | 0 | 6 |
| E47N | 0 | 8 | 6 | 6 | 8 | 5 | 7 | 0 | 7 | 5 | 5 | 6 | 5 | 0 | 0 | 0 | 0 | 0 | 0 | 5 |
| L50F | 0 | 9 | 0 | 8 | 7 | 0 | 8 | 0 | 5 | 4 | 0 | 2 | 3 | 0 | 4 | 0 | 5 | 0 | 0 | 4 |
| L89F | 0 | 7 | 4 | 4 | 0 | 0 | 6 | 0 | 0 | 0 | 0 | 5 | 0 | 0 | 0 | 0 | 0 | 0 | 0 | 0 |
| K90R | 6 | 12 | 9 | 11 | 9 | 7 | 9 | 0 | 7 | 7 | 6 | 9 | 7 | 8 | 8 | 8 | 7 | 0 | 0 | 9 |
| P108S | 0 | 13 | 5 | 9 | 8 | 7 | 8 | 0 | 0 | 7 | 6 | 9 | 8 | 7 | 7 | 5 | 7 | 0 | 0 | 7 |
| P132H | 0 | 15 | 13 | 13 | 14 | 10 | 11 | 0 | 0 | 10 | 0 | 11 | 0 | 0 | 0 | 0 | 0 | 0 | 0 | 9 |
| A260V | 8 | 16 | 13 | 11 | 22 | 8 | 10 | 7 | 11 | 8 | 9 | 11 | 10 | 8 | 9 | 8 | 9 | 5 | 0 | 8 |
| G283S | 6 | 9 | 6 | 7 | 5 | 4 | 6 | 0 | 6 | 5 | 6 | 6 | 5 | 6 | 5 | 2 | 5 | 4 | 0 | 6 |
| S284G | 5 | 12 | 8 | 9 | 11 | 7 | 9 | 4 | 5 | 7 | 5 | 8 | 6 | 6 | 6 | 5 | 7 | 0 | 0 | 6 |

| Variant | L-Agb | L-Arg(Z)2 | L-hArg | L-His(3-Bom) | L-Phe(NH2) | L-Phe(guan) | L-Trp(Me) | L-Dht | L-Asp(O-Me) | L-Asp(O-CHx) | L-Asp(O-Bzl) | L-Asp(OAll) | L-Glu(O-Me) | L-Glu(O-CHx) | L-Glu(O-Bzl) | L-Glu(All) | L-Aad | L-Phe(2-F) | L-Phe(3-F) | L-Phe(4-F) |
|---|---|---|---|---|---|---|---|---|---|---|---|---|---|---|---|---|---|---|---|---|
| WT | 11 | 0 | 3 | 4 | 5 | 3 | 3 | 9 | 5 | 4 | 6 | 6 | 5 | 5 | 4 | 4 | 0 | 4 | 0 | 4 |
| E47K | 15 | 0 | 0 | 0 | 0 | 8 | 10 | 12 | 11 | 11 | 12 | 13 | 11 | 13 | 12 | 11 | 8 | 10 | 10 | 11 |
| E47N | 15 | 0 | 5 | 2 | 5 | 4 | 6 | 8 | 6 | 5 | 7 | 7 | 4 | 8 | 6 | 4 | 1 | 4 | 4 | 4 |
| L50F | 9 | 0 | 3 | 0 | 7 | 3 | 8 | 17 | 0 | 9 | 0 | 0 | 0 | 12 | 5 | 6 | 0 | 0 | 6 | 7 |
| L89F | 11 | 0 | 3 | 0 | 4 | 4 | 7 | 9 | 8 | 0 | 4 | 7 | 7 | 0 | 0 | 6 | 0 | 0 | 0 | 0 |
| K90R | 18 | 3 | 9 | 0 | 7 | 5 | 9 | 10 | 5 | 7 | 6 | 6 | 6 | 8 | 7 | 7 | 0 | 4 | 4 | 6 |
| P108S | 18 | 0 | 8 | 0 | 7 | 3 | 9 | 10 | 0 | 0 | 0 | 0 | 0 | 8 | 0 | 0 | 0 | 0 | 0 | 0 |
| P132H | 21 | 0 | 11 | 0 | 8 | 4 | 6 | 10 | 5 | 6 | 6 | 7 | 5 | 7 | 4 | 3 | 0 | 2 | 6 | 6 |
| A260V | 21 | 5 | 11 | 5 | 9 | 9 | 8 | 12 | 9 | 8 | 8 | 9 | 8 | 8 | 11 | 10 | 0 | 0 | 4 | 7 |
| G283S | 16 | 4 | 4 | 2 | 7 | 3 | 6 | 13 | 4 | 9 | 6 | 6 | 3 | 6 | 6 | 10 | 0 | 5 | 6 | 6 |
| S284G | 17 | 7 | 8 | 4 | 8 | 6 | 8 | 14 | 7 | 8 | 6 | 7 | 8 | 9 | 9 | 5 | 0 | 8 | 8 | 8 |

| Variant | L-Phe(3,4-F2) | L-Phe(F5) | L-Phe(2-Cl) | L-Phe(3-Cl) | L-Phe(4-Cl) | L-Phe(3,4-Cl2) | L-Phe(4-Br) | L-Phe(3-I) | L-Phe(4-I) | L-Phe(4-Me) | L-3-Pal | L-4-Pal | L-Ala(2-th) | L-Ala(Bth) | L-Abu | L-Abu(Bth) | L-Ser(Ac) | L-Ser(Bzl) | L-hSer | L-Thr(Bzl) |
|---|---|---|---|---|---|---|---|---|---|---|---|---|---|---|---|---|---|---|---|---|
| WT | 0 | 2 | 6 | 0 | 3 | 0 | 3 | 3 | 2 | 3 | 3 | 5 | 5 | 0 | 0 | 0 | 6 | 3 | 4 | 3 |
| E47K | 8 | 8 | 14 | 0 | 9 | 7 | 7 | 5 | 8 | 7 | 9 | 14 | 11 | 6 | 8 | 6 | 13 | 12 | 7 | 10 |
| E47N | 0 | 0 | 4 | 0 | 0 | 0 | 0 | 0 | 0 | 3 | 0 | 6 | 6 | 0 | 0 | 0 | 5 | 8 | 2 | 7 |
| L50F | 0 | 0 | 5 | 0 | 0 | 0 | 0 | 0 | 0 | 4 | 0 | 8 | 7 | 0 | 0 | 0 | 0 | 0 | 0 | 0 |
| L89F | 0 | 0 | 7 | 0 | 0 | 0 | 0 | 0 | 0 | 0 | 0 | 5 | 5 | 0 | 0 | 0 | 6 | 4 | 0 | 0 |
| K90R | 0 | 0 | 7 | 0 | 0 | 0 | 0 | 0 | 1 | 2 | 4 | 7 | 5 | 2 | 0 | 0 | 0 | 5 | 0 | 0 |
| P108S | 0 | 0 | 0 | 0 | 0 | 0 | 0 | 0 | 0 | 0 | 0 | 4 | 7 | 0 | 0 | 0 | 0 | 0 | 0 | 0 |
| P132H | 0 | 0 | 8 | 0 | 0 | 0 | 0 | 0 | 0 | 0 | 0 | 8 | 6 | 0 | 4 | 0 | 0 | 7 | 0 | 0 |
| A260V | 5 | 0 | 8 | 0 | 5 | 0 | 4 | 4 | 6 | 4 | 5 | 9 | 8 | 4 | 0 | 0 | 7 | 6 | 4 | 0 |
| G283S | 3 | 0 | 6 | 0 | 0 | 0 | 9 | 0 | 0 | 0 | 0 | 6 | 4 | 0 | 0 | 0 | 0 | 8 | 7 | 6 |
| S284G | 0 | 7 | 9 | 4 | 7 | 5 | 6 | 2 | 5 | 6 | 4 | 9 | 9 | 5 | 5 | 0 | 0 | 8 | 4 | 8 |

| Variant | L-Cys(Bzl) | L-Cys(MeBzl) | L-Cys(4-MeOBzl) | L-Met | L-Met(O) | L-Met(O)2 | L-Nle(O-Bzl) | L-Phg | L-hPhe | L-Chg | L-Cha | L-hCha | L-Igl | L-1-Nal | L-2-Nal | L-Bip | L-Bpa | L-2-Aoc | L-hLeu | L-Tle |
|---|---|---|---|---|---|---|---|---|---|---|---|---|---|---|---|---|---|---|---|---|
| WT | 5 | 2 | 3 | 15 | 2 | 4 | 0 | 8 | 7 | 11 | 10 | 0 | 5 | 3 | 3 | 0 | 0 | 2 | 17 | 8 |
| E47K | 7 | 6 | 10 | 20 | 8 | 8 | 8 | 14 | 12 | 15 | 13 | 8 | 11 | 10 | 9 | 6 | 8 | 8 | 18 | 17 |
| E47N | 6 | 0 | 5 | 14 | 0 | 0 | 0 | 10 | 7 | 11 | 9 | 0 | 6 | 7 | 5 | 0 | 0 | 0 | 15 | 6 |
| L50F | 0 | 0 | 0 | 14 | 0 | 0 | 0 | 0 | 0 | 0 | 0 | 0 | 0 | 15 | 0 | 5 | 7 | 0 | 0 | 10 |
| L89F | 0 | 0 | 8 | 16 | 0 | 0 | 0 | 6 | 8 | 11 | 10 | 4 | 6 | 5 | 0 | 0 | 4 | 5 | 15 | 8 |
| K90R | 0 | 0 | 5 | 15 | 0 | 3 | 4 | 8 | 6 | 11 | 8 | 2 | 7 | 6 | 4 | 4 | 4 | 0 | 13 | 8 |
| P108S | 0 | 0 | 6 | 14 | 0 | 0 | 0 | 9 | 0 | 10 | 0 | 0 | 0 | 4 | 0 | 0 | 0 | 0 | 13 | 7 |
| P132H | 5 | 0 | 6 | 15 | 0 | 2 | 3 | 8 | 2 | 8 | 5 | 0 | 0 | 7 | 0 | 0 | 0 | 0 | 12 | 4 |
| A260V | 7 | 0 | 6 | 15 | 0 | 6 | 6 | 10 | 13 | 11 | 8 | 0 | 0 | 0 | 7 | 0 | 4 | 7 | 12 | 8 |
| G283S | 0 | 0 | 4 | 11 | 0 | 0 | 0 | 6 | 4 | 7 | 3 | 0 | 6 | 9 | 0 | 4 | 0 | 0 | 12 | 7 |
| S284G | 6 | 4 | 9 | 16 | 0 | 6 | 0 | 9 | 5 | 11 | 6 | 9 | 7 | 6 | 4 | 5 | 6 | 0 | 13 | 9 |

**Figure S6.** Substrate specificity profile of SARS-CoV-2 M^pro presented as heat maps using substrate library with unnatural amino acids assessing the P2 position with Ac-Mix-Mix-P2-Gln-ACC.

| Variant | L-Arg(NO2) | L-Tyr(Me) | L-Tyr(2,6-Cl2-Bzl) | L-Tyr(Bzl) | L-Tyr(2-Br-Z) | L-hTyr | L-hTyr(Me) | L-Nva | L-His(Bzl) | L-2-Abz | L-3-Abz | L-4-Abz | Inp | L-Gla | L-Phe(4-NO2) | Aib | L-hSer(Bzl) | TmbGly | L-Nle |
|---|---|---|---|---|---|---|---|---|---|---|---|---|---|---|---|---|---|---|---|
| WT | 3 | 4 | 0 | 0 | 6 | 4 | 4 | 12 | 8 | 7 | 10 | 5 | 5 | 5 | 11 | 4 | 5 | 9 | 19 |
| E47K | 9 | 10 | 11 | 7 | 14 | 11 | 10 | 19 | 15 | 16 | 23 | 15 | 11 | 13 | 20 | 11 | 12 | 17 | 19 |
| E47N | 0 | 5 | 4 | 0 | 8 | 5 | 5 | 12 | 7 | 8 | 14 | 8 | 5 | 5 | 13 | 6 | 0 | 8 | 20 |
| L50F | 0 | 4 | 0 | 6 | 12 | 8 | 0 | 13 | 12 | 0 | 8 | 0 | 0 | 0 | 0 | 0 | 8 | 7 | 17 |
| L89F | 5 | 0 | 0 | 0 | 8 | 5 | 5 | 13 | 10 | 11 | 13 | 4 | 0 | 0 | 12 | 0 | 0 | 0 | 20 |
| K90R | 4 | 6 | 6 | 2 | 10 | 6 | 5 | 13 | 15 | 11 | 17 | 9 | 2 | 4 | 12 | 6 | 7 | 7 | 23 |
| P108S | 0 | 6 | 4 | 0 | 9 | 6 | 0 | 11 | 12 | 8 | 12 | 11 | 4 | 0 | 14 | 0 | 7 | 7 | 23 |
| P132H | 2 | 6 | 4 | 0 | 9 | 5 | 5 | 12 | 9 | 8 | 13 | 6 | 4 | 0 | 12 | 5 | 7 | 11 | 25 |
| A260V | 7 | 3 | 7 | 6 | 8 | 5 | 6 | 13 | 11 | 14 | 15 | 12 | 9 | 9 | 18 | 8 | 6 | 12 | 22 |
| G283S | 0 | 5 | 0 | 0 | 9 | 6 | 0 | 9 | 9 | 7 | 12 | 10 | 5 | 0 | 13 | 0 | 7 | 5 | 20 |
| S284G | 3 | 5 | 6 | 0 | 9 | 7 | 0 | 14 | 11 | 11 | 14 | 12 | 7 | 0 | 16 | 6 | 8 | 10 | 24 |

Ac-Mix-P3-Mix-Gln-ACC

| Variants | D-Ala | D-Arg | D-Asn | D-Asp | D-Gln | D-Glu | D-His | D-Leu | D-Lys | D-Phe | D-Pro | D-Ser | D-Phg | D-Thr | D-Trp | D-Tyr | D-Val | D-hPhe | B-Ala | L-Aze |
|---|---|---|---|---|---|---|---|---|---|---|---|---|---|---|---|---|---|---|---|---|
| WT | 0 | 0 | 0 | 0 | 0 | 0 | 0 | 17 | 0 | 47 | 0 | 0 | 31 | 0 | 29 | 71 | 9 | 8 | 0 | 0 |
| E47K | 0 | 0 | 0 | 0 | 0 | 0 | 0 | 18 | 0 | 39 | 0 | 0 | 34 | 0 | 28 | 62 | 8 | 8 | 0 | 0 |
| E47N | 0 | 0 | 0 | 0 | 0 | 0 | 0 | 18 | 0 | 43 | 0 | 0 | 36 | 0 | 32 | 74 | 8 | 9 | 0 | 0 |
| L50F | 0 | 0 | 0 | 0 | 0 | 0 | 0 | 9 | 0 | 22 | 0 | 0 | 19 | 0 | 15 | 30 | 0 | 7 | 0 | 0 |
| L89F | 0 | 0 | 0 | 0 | 0 | 0 | 0 | 12 | 0 | 41 | 0 | 0 | 40 | 0 | 33 | 68 | 0 | 0 | 0 | 0 |
| K90R | 0 | 0 | 0 | 0 | 0 | 0 | 0 | 26 | 0 | 48 | 0 | 0 | 42 | 0 | 32 | 78 | 16 | 19 | 0 | 0 |
| P108S | 0 | 0 | 0 | 0 | 0 | 0 | 0 | 16 | 0 | 38 | 0 | 0 | 33 | 0 | 26 | 63 | 0 | 0 | 0 | 0 |
| P132H | 0 | 0 | 0 | 0 | 0 | 0 | 0 | 20 | 0 | 41 | 0 | 0 | 33 | 0 | 26 | 75 | 0 | 4 | 0 | 0 |
| A260V | 0 | 0 | 0 | 0 | 0 | 0 | 0 | 10 | 0 | 33 | 0 | 0 | 24 | 0 | 17 | 55 | 0 | 0 | 0 | 0 |
| G283S | 0 | 0 | 0 | 0 | 0 | 0 | 0 | 11 | 0 | 37 | 0 | 0 | 29 | 0 | 24 | 65 | 0 | 0 | 0 | 0 |
| S284G | 0 | 0 | 0 | 0 | 0 | 0 | 0 | 14 | 0 | 38 | 0 | 0 | 32 | 0 | 21 | 64 | 0 | 15 | 0 | 0 |

| Variant | L-Hyp | L-Hyp(Bzl) | L-Thz | L-Oic | L-Idc | L-Pip | L-Tic | AC5C | dhAbu | dhLeu | L-Dap | L-Dab | L-Dab(Z) | L-Cit | L-hCit | L-Orn | L-Lys(TFA) | L-Lys(Ac) | L-Lys(2-ClZ) | L-Agp |
|---|---|---|---|---|---|---|---|---|---|---|---|---|---|---|---|---|---|---|---|---|
| WT | 0 | 0 | 0 | 0 | 12 | 0 | 0 | 0 | 0 | 10 | 17 | 43 | 11 | 15 | 9 | 43 | 15 | 16 | 14 | 32 |
| E47K | 0 | 0 | 0 | 0 | 0 | 0 | 0 | 0 | 0 | 9 | 14 | 41 | 16 | 14 | 14 | 41 | 19 | 14 | 0 | 33 |
| E47N | 0 | 0 | 0 | 0 | 11 | 0 | 0 | 0 | 0 | 10 | 17 | 51 | 21 | 18 | 12 | 48 | 17 | 14 | 0 | 37 |
| L50F | 0 | 0 | 0 | 0 | 0 | 0 | 0 | 0 | 0 | 0 | 11 | 47 | 20 | 16 | 20 | 34 | 15 | 15 | 0 | 31 |
| L89F | 0 | 0 | 0 | 0 | 5 | 0 | 0 | 0 | 0 | 7 | 17 | 57 | 14 | 12 | 0 | 53 | 13 | 12 | 20 | 43 |
| K90R | 0 | 0 | 0 | 0 | 0 | 0 | 0 | 0 | 0 | 17 | 22 | 56 | 18 | 25 | 22 | 58 | 25 | 24 | 12 | 46 |
| P108S | 0 | 0 | 0 | 0 | 0 | 0 | 0 | 0 | 0 | 9 | 11 | 47 | 0 | 15 | 0 | 41 | 16 | 13 | 0 | 36 |
| P132H | 0 | 0 | 0 | 0 | 0 | 0 | 0 | 0 | 0 | 12 | 15 | 46 | 0 | 14 | 10 | 45 | 16 | 14 | 23 | 38 |
| A260V | 0 | 0 | 0 | 0 | 6 | 0 | 0 | 0 | 0 | 0 | 8 | 32 | 8 | 12 | 0 | 34 | 12 | 11 | 26 | 28 |
| G283S | 0 | 0 | 0 | 0 | 0 | 0 | 0 | 0 | 0 | 8 | 11 | 43 | 17 | 7 | 0 | 44 | 13 | 14 | 0 | 34 |
| S284G | 0 | 0 | 0 | 0 | 0 | 0 | 0 | 0 | 0 | 9 | 16 | 44 | 9 | 13 | 13 | 46 | 14 | 7 | 11 | 31 |

| Variant | L-Agb | L-Arg(Z)2 | L-hArg | L-His(3-Bom) | L-Phe(NH2) | L-Phe(guan) | L-Trp(Me) | L-Dht | L-Asp(O-Me) | L-Asp(O-CHx) | L-Asp(O-Bzl) | L-Asp(OAll) | L-Glu(O-Me) | L-Glu(O-CHx) | L-Glu(O-Bzl) | L-Glu(All) | L-Aad | L-Phe(2-F) | L-Phe(3-F) | L-Phe(4-F) |
|---|---|---|---|---|---|---|---|---|---|---|---|---|---|---|---|---|---|---|---|---|
| WT | 14 | 0 | 36 | 4 | 9 | 26 | 8 | 38 | 0 | 0 | 0 | 0 | 9 | 9 | 10 | 6 | 0 | 0 | 0 | 0 |
| E47K | 15 | 0 | 36 | 0 | 13 | 18 | 0 | 30 | 0 | 0 | 0 | 0 | 0 | 0 | 0 | 0 | 0 | 0 | 0 | 0 |
| E47N | 16 | 0 | 41 | 8 | 11 | 27 | 5 | 36 | 0 | 0 | 0 | 0 | 12 | 0 | 0 | 0 | 0 | 0 | 0 | 0 |
| L50F | 12 | 0 | 48 | 10 | 15 | 26 | 10 | 40 | 0 | 7 | 0 | 0 | 19 | 8 | 9 | 14 | 0 | 0 | 0 | 0 |
| L89F | 13 | 0 | 36 | 0 | 9 | 31 | 3 | 39 | 0 | 0 | 0 | 0 | 11 | 0 | 0 | 0 | 0 | 0 | 0 | 0 |
| K90R | 27 | 0 | 53 | 0 | 19 | 25 | 0 | 35 | 0 | 0 | 0 | 0 | 0 | 0 | 0 | 0 | 0 | 0 | 0 | 0 |
| P108S | 18 | 0 | 36 | 0 | 14 | 25 | 0 | 32 | 0 | 0 | 0 | 0 | 0 | 0 | 0 | 0 | 0 | 0 | 0 | 0 |
| P132H | 14 | 0 | 38 | 0 | 16 | 27 | 9 | 37 | 0 | 0 | 0 | 0 | 0 | 0 | 0 | 0 | 0 | 0 | 0 | 0 |
| A260V | 10 | 0 | 32 | 10 | 9 | 21 | 0 | 29 | 0 | 0 | 0 | 0 | 11 | 0 | 0 | 0 | 0 | 0 | 0 | 0 |
| G283S | 12 | 0 | 29 | 0 | 9 | 22 | 0 | 35 | 0 | 0 | 0 | 0 | 0 | 8 | 0 | 0 | 0 | 0 | 0 | 0 |
| S284G | 16 | 0 | 40 | 0 | 10 | 25 | 0 | 36 | 0 | 0 | 0 | 0 | 14 | 0 | 0 | 11 | 0 | 0 | 0 | 0 |

**Figure S6 continued.** Substrate specificity profile of SARS-CoV-2 $M^{pro}$ presented as heat maps using substrate library with unnatural amino acids assessing the P2 and P3 positions with Ac-Mix-Mix-P2-Gln-ACC and Ac-Mix-P3-Mix-Gln-ACC.

| Variant | L-Phe(3,4-F$_2$) | L-Phe(F$_5$) | L-Phe(2-Cl) | L-Phe(3-Cl) | L-Phe(4-Cl) | L-Phe(3,4-Cl$_2$) | L-Phe(4-Br) | L-Phe(3-I) | L-Phe(4-I) | L-Phe(4-Me) | L-3-Pal | L-4-Pal | L-Ala(2-th) | L-Ala(Bth) | L-Abu | L-Abu(Bth) | L-Ser(Ac) | L-Ser(Bzl) | L-hSer | L-Thr(Bzl) |
|---|---|---|---|---|---|---|---|---|---|---|---|---|---|---|---|---|---|---|---|---|
| WT | 0 | 0 | 0 | 0 | 0 | 0 | 0 | 0 | 0 | 0 | 8 | 9 | 0 | 0 | 10 | 0 | 0 | 0 | 9 | 7 |
| E47K | 0 | 0 | 0 | 0 | 0 | 0 | 0 | 0 | 0 | 0 | 0 | 0 | 0 | 0 | 0 | 0 | 0 | 0 | 0 | 0 |
| E47N | 0 | 12 | 0 | 0 | 0 | 0 | 0 | 0 | 0 | 0 | 12 | 12 | 0 | 0 | 13 | 0 | 0 | 0 | 12 | 0 |
| L50F | 0 | 8 | 0 | 0 | 0 | 0 | 0 | 0 | 0 | 0 | 14 | 16 | 0 | 0 | 16 | 1 | 0 | 0 | 13 | 6 |
| L89F | 0 | 4 | 0 | 0 | 0 | 0 | 0 | 0 | 0 | 0 | 10 | 12 | 0 | 0 | 10 | 0 | 0 | 0 | 11 | 0 |
| K90R | 0 | 0 | 0 | 0 | 0 | 0 | 0 | 0 | 0 | 0 | 0 | 0 | 0 | 0 | 0 | 0 | 0 | 0 | 0 | 0 |
| P108S | 0 | 0 | 0 | 0 | 0 | 0 | 0 | 0 | 0 | 0 | 0 | 0 | 0 | 0 | 0 | 0 | 0 | 0 | 0 | 0 |
| P132H | 0 | 0 | 0 | 0 | 0 | 0 | 0 | 0 | 0 | 0 | 0 | 0 | 0 | 0 | 0 | 0 | 0 | 0 | 0 | 0 |
| A260V | 0 | 0 | 0 | 0 | 0 | 0 | 0 | 0 | 0 | 0 | 0 | 0 | 0 | 0 | 0 | 0 | 0 | 0 | 12 | 0 |
| G283S | 0 | 0 | 0 | 0 | 0 | 0 | 0 | 0 | 0 | 0 | 10 | 12 | 0 | 0 | 0 | 0 | 0 | 0 | 0 | 0 |
| S284G | 0 | 0 | 0 | 0 | 0 | 0 | 0 | 0 | 0 | 0 | 11 | 17 | 0 | 0 | 11 | 0 | 0 | 0 | 10 | 0 |

| Variant | L-Cys(Bzl) | L-Cys(MeBzl) | L-Cys(4-MeOBzl) | L-Met | L-Met(O) | L-Met(O)$_2$ | L-Nle(O-Bzl) | L-Phg | L-hPhe | L-Chg | L-Cha | L-Igl | L-1-Nal | L-2-Nal | L-Bip | L-Bpa | L-2-Aoc | L-Tle | L-Arg(NO$_2$) | L-Tyr(Me) |
|---|---|---|---|---|---|---|---|---|---|---|---|---|---|---|---|---|---|---|---|---|
| WT | 0 | 5 | 11 | 15 | 18 | 28 | 0 | 23 | 3 | 8 | 0 | 0 | 7 | 0 | 0 | 5 | 0 | 100 | 23 | 0 |
| E47K | 0 | 0 | 13 | 12 | 8 | 27 | 0 | 26 | 0 | 0 | 0 | 0 | 12 | 0 | 0 | 0 | 0 | 100 | 19 | 0 |
| E47N | 0 | 7 | 17 | 20 | 24 | 36 | 0 | 41 | 7 | 15 | 0 | 0 | 11 | 0 | 0 | 7 | 0 | 100 | 28 | 12 |
| L50F | 0 | 4 | 25 | 25 | 25 | 39 | 11 | 29 | 18 | 14 | 11 | 0 | 17 | 0 | 0 | 5 | 7 | 100 | 28 | 10 |
| L89F | 0 | 0 | 16 | 21 | 22 | 36 | 0 | 28 | 10 | 10 | 3 | 0 | 0 | 0 | 0 | 0 | 0 | 100 | 30 | 0 |
| K90R | 0 | 0 | 26 | 15 | 25 | 31 | 0 | 29 | 0 | 0 | 0 | 0 | 0 | 0 | 0 | 0 | 0 | 100 | 23 | 0 |
| P108S | 0 | 0 | 0 | 20 | 17 | 36 | 0 | 19 | 0 | 0 | 0 | 0 | 0 | 0 | 0 | 0 | 0 | 100 | 24 | 0 |
| P132H | 0 | 0 | 19 | 21 | 21 | 36 | 0 | 20 | 0 | 0 | 0 | 0 | 0 | 0 | 0 | 0 | 0 | 100 | 27 | 0 |
| A260V | 0 | 0 | 12 | 13 | 18 | 26 | 0 | 15 | 0 | 12 | 0 | 0 | 17 | 0 | 0 | 0 | 0 | 100 | 22 | 0 |
| G283S | 0 | 0 | 15 | 14 | 19 | 35 | 0 | 18 | 11 | 0 | 0 | 0 | 0 | 0 | 0 | 0 | 0 | 100 | 21 | 0 |
| S284G | 0 | 0 | 23 | 18 | 25 | 38 | 0 | 23 | 0 | 16 | 0 | 0 | 9 | 8 | 0 | 0 | 0 | 100 | 25 | 6 |

| Variant | L-Tyr(2,6-Cl$_2$-Bzl) | L-Tyr(Bzl) | L-Tyr(2-Br-Z) | L-hTyr | L-hTyr(Me) | L-Nva | L-His(Bzl) | L-2-Abz | L-4-Abz | Inp | L-Gla | L-Phe(4-NO2) | Aib | L-hSer(Bzl) | L-Nle |
|---|---|---|---|---|---|---|---|---|---|---|---|---|---|---|---|
| WT | 0 | 5 | 11 | 12 | 0 | 9 | 4 | 0 | 0 | 0 | 0 | 0 | 0 | 0 | 9 |
| E47K | 0 | 0 | 0 | 14 | 0 | 4 | 0 | 0 | 0 | 0 | 0 | 0 | 0 | 0 | 0 |
| E47N | 0 | 0 | 18 | 24 | 9 | 17 | 5 | 0 | 0 | 0 | 0 | 0 | 0 | 0 | 6 |
| L50F | 0 | 9 | 18 | 21 | 11 | 17 | 8 | 0 | 0 | 0 | 0 | 0 | 0 | 8 | 0 |
| L89F | 0 | 0 | 15 | 19 | 10 | 14 | 0 | 0 | 0 | 0 | 0 | 0 | 0 | 0 | 5 |
| K90R | 0 | 0 | 0 | 22 | 22 | 0 | 0 | 0 | 0 | 0 | 0 | 0 | 0 | 0 | 17 |
| P108S | 0 | 0 | 13 | 20 | 0 | 0 | 0 | 0 | 0 | 0 | 0 | 0 | 0 | 0 | 0 |
| P132H | 0 | 0 | 15 | 13 | 0 | 0 | 0 | 0 | 0 | 0 | 0 | 0 | 0 | 0 | 0 |
| A260V | 0 | 6 | 8 | 6 | 9 | 9 | 0 | 0 | 0 | 0 | 0 | 0 | 0 | 0 | 0 |
| G283S | 0 | 0 | 12 | 14 | 0 | 17 | 5 | 0 | 0 | 0 | 0 | 0 | 0 | 0 | 0 |
| S284G | 0 | 8 | 13 | 17 | 0 | 16 | 0 | 0 | 0 | 0 | 0 | 0 | 0 | 0 | 8 |

**Figure S6 continued.** Substrate specificity profile of SARS-CoV-2 M$^{pro}$ presented as heat maps using substrate library with unnatural amino acids assessing the P3 position with Ac-Mix-P3-Mix-Gln-ACC.

Ac-P4-Mix-Mix-Gln-ACC

| Variant | D-Ala | D-Arg | D-Asn | D-Asp | D-Gln | D-Glu | D-His | D-Leu | D-Lys | D-Phe | D-Pro | D-Ser | D-Phg | D-Thr | D-Trp | D-Tyr | D-Val | D-hPhe | B-Ala | L-Aze |
|---|---|---|---|---|---|---|---|---|---|---|---|---|---|---|---|---|---|---|---|---|
| WT | 0 | 0 | 0 | 0 | 0 | 0 | 0 | 0 | 0 | 0 | 0 | 0 | 19 | 0 | 0 | 0 | 0 | 0 | 18 | 32 |
| E47K | 0 | 0 | 0 | 0 | 0 | 0 | 0 | 0 | 0 | 0 | 0 | 0 | 13 | 0 | 0 | 0 | 0 | 0 | 0 | 17 |
| E47N | 0 | 0 | 0 | 0 | 0 | 0 | 0 | 0 | 0 | 0 | 0 | 0 | 0 | 0 | 0 | 0 | 0 | 0 | 0 | 0 |
| L50F | 0 | 0 | 0 | 0 | 0 | 0 | 0 | 0 | 0 | 0 | 0 | 0 | 9 | 0 | 0 | 0 | 0 | 0 | 0 | 0 |
| L89F | 0 | 0 | 0 | 0 | 0 | 0 | 0 | 0 | 0 | 0 | 0 | 0 | 0 | 0 | 0 | 0 | 0 | 0 | 0 | 27 |
| K90R | 0 | 0 | 0 | 0 | 0 | 0 | 0 | 0 | 0 | 0 | 0 | 0 | 0 | 0 | 0 | 0 | 0 | 0 | 0 | 0 |
| P108S | 0 | 0 | 0 | 0 | 0 | 0 | 0 | 0 | 0 | 0 | 0 | 0 | 0 | 0 | 0 | 0 | 0 | 0 | 0 | 0 |
| P132H | 0 | 0 | 0 | 0 | 0 | 0 | 0 | 0 | 0 | 0 | 0 | 0 | 0 | 0 | 0 | 0 | 0 | 0 | 0 | 21 |
| A260V | 0 | 0 | 0 | 0 | 0 | 0 | 0 | 0 | 0 | 0 | 0 | 0 | 0 | 0 | 0 | 0 | 0 | 0 | 0 | 0 |
| G283S | 0 | 0 | 0 | 0 | 0 | 0 | 0 | 0 | 0 | 0 | 0 | 0 | 17 | 0 | 0 | 0 | 0 | 0 | 0 | 26 |
| S284G | 0 | 0 | 0 | 0 | 0 | 0 | 0 | 0 | 0 | 0 | 0 | 0 | 9 | 0 | 0 | 0 | 0 | 0 | 0 | 27 |

| Variant | L-Hyp | L-Hyp(Bzl) | L-Thz | L-Oic | L-Idc | L-Pip | L-Tic | AC5C | dhAbu | dhLeu | L-Dap | L-Dab | L-Dab(Z) | L-Cit | L-hCit | L-Orn | L-Lys(TFA) | L-Lys(Ac) | L-Lys(2-ClZ) | L-Agp |
|---|---|---|---|---|---|---|---|---|---|---|---|---|---|---|---|---|---|---|---|---|
| WT | 9 | 19 | 99 | 0 | 0 | 8 | 7 | 0 | 38 | 8 | 0 | 0 | 0 | 0 | 0 | 0 | 0 | 0 | 0 | 0 |
| E47K | 0 | 20 | 63 | 0 | 0 | 0 | 0 | 0 | 26 | 0 | 0 | 0 | 0 | 0 | 0 | 0 | 0 | 0 | 0 | 0 |
| E47N | 0 | 0 | 66 | 0 | 0 | 0 | 0 | 0 | 24 | 0 | 0 | 0 | 0 | 0 | 0 | 0 | 0 | 0 | 0 | 0 |
| L50F | 0 | 0 | 65 | 0 | 0 | 0 | 0 | 0 | 0 | 0 | 0 | 0 | 0 | 0 | 0 | 0 | 0 | 0 | 0 | 0 |
| L89F | 0 | 22 | 73 | 0 | 0 | 0 | 0 | 0 | 30 | 0 | 0 | 0 | 0 | 0 | 0 | 0 | 0 | 0 | 0 | 0 |
| K90R | 0 | 0 | 66 | 0 | 0 | 0 | 0 | 0 | 0 | 0 | 0 | 0 | 0 | 0 | 0 | 0 | 0 | 0 | 0 | 0 |
| P108S | 0 | 17 | 64 | 0 | 0 | 0 | 0 | 0 | 19 | 0 | 0 | 0 | 0 | 0 | 0 | 0 | 0 | 0 | 0 | 0 |
| P132H | 0 | 19 | 63 | 0 | 0 | 0 | 0 | 0 | 25 | 0 | 0 | 0 | 0 | 0 | 0 | 0 | 0 | 0 | 0 | 0 |
| A260V | 0 | 0 | 61 | 0 | 0 | 0 | 0 | 0 | 15 | 0 | 0 | 0 | 0 | 0 | 0 | 0 | 0 | 0 | 0 | 0 |
| G283S | 0 | 8 | 63 | 0 | 11 | 0 | 0 | 0 | 24 | 10 | 0 | 0 | 0 | 0 | 0 | 0 | 0 | 0 | 0 | 24 |
| S284G | 0 | 10 | 69 | 0 | 0 | 0 | 0 | 0 | 15 | 0 | 0 | 0 | 0 | 0 | 0 | 0 | 0 | 0 | 0 | 0 |

| Variant | L-Agb | L-Arg(Z)2 | L-hArg | L-His(3-Bom) | L-Phe(NH2) | L-Phe(guan) | L-Trp(Me) | L-Dht | L-Asp(O-Me) | L-Asp(O-CHx) | L-Asp(O-Bzl) | L-Asp(OAll) | L-Glu(O-Me) | L-Glu(O-CHx) | L-Glu(O-Bzl) | L-Glu(All) | L-Aad | L-Phe(2-F) | L-Phe(3-F) | L-Phe(4-F) |
|---|---|---|---|---|---|---|---|---|---|---|---|---|---|---|---|---|---|---|---|---|
| WT | 25 | 0 | 0 | 0 | 0 | 0 | 0 | 0 | 0 | 0 | 0 | 0 | 0 | 0 | 0 | 10 | 0 | 0 | 0 | 0 |
| E47K | 15 | 0 | 0 | 0 | 0 | 0 | 0 | 0 | 0 | 0 | 0 | 0 | 0 | 0 | 0 | 0 | 0 | 0 | 0 | 0 |
| E47N | 0 | 0 | 0 | 0 | 0 | 0 | 0 | 17 | 0 | 0 | 0 | 0 | 0 | 0 | 21 | 15 | 0 | 0 | 0 | 0 |
| L50F | 0 | 0 | 0 | 0 | 0 | 0 | 0 | 0 | 0 | 0 | 0 | 0 | 0 | 0 | 0 | 0 | 0 | 0 | 0 | 0 |
| L89F | 22 | 0 | 0 | 0 | 0 | 0 | 0 | 0 | 0 | 0 | 0 | 0 | 0 | 0 | 0 | 0 | 0 | 0 | 0 | 0 |
| K90R | 0 | 0 | 0 | 0 | 0 | 0 | 0 | 0 | 0 | 0 | 0 | 0 | 0 | 0 | 0 | 0 | 0 | 0 | 0 | 0 |
| P108S | 22 | 0 | 0 | 0 | 0 | 0 | 0 | 0 | 0 | 0 | 0 | 0 | 0 | 0 | 0 | 0 | 0 | 0 | 0 | 0 |
| P132H | 0 | 0 | 0 | 0 | 0 | 0 | 0 | 0 | 0 | 0 | 0 | 0 | 0 | 0 | 0 | 0 | 0 | 0 | 0 | 0 |
| A260V | 0 | 0 | 0 | 0 | 0 | 0 | 0 | 8 | 0 | 0 | 0 | 0 | 0 | 0 | 0 | 0 | 0 | 0 | 0 | 0 |
| G283S | 0 | 0 | 0 | 0 | 0 | 0 | 0 | 24 | 0 | 0 | 0 | 0 | 0 | 0 | 0 | 0 | 0 | 0 | 0 | 0 |
| S284G | 15 | 0 | 0 | 0 | 0 | 0 | 0 | 20 | 0 | 0 | 0 | 0 | 0 | 0 | 0 | 0 | 0 | 0 | 0 | 0 |

**Figure S6 continued.** Substrate specificity profile of SARS-CoV-2 M$^{pro}$ presented as heat maps using substrate library with unnatural amino acids assessing the P4 position with Ac-P4-Mix-Mix-Gln-ACC.

| Variant | L-Phe(3,4-F$_2$) | L-Phe(F$_5$) | L-Phe(2-Cl) | L-Phe(3-Cl) | L-Phe(4-Cl) | L-Phe(3,4-Cl$_2$) | L-Phe(4-Br) | L-Phe(3-I) | L-Phe(4-I) | L-Phe(4-Me) | L-3-Pal | L-4-Pal | L-Ala(2-th) | L-Ala(Bth) | L-Abu | L-Abu(Bth) | L-Ser(Ac) | L-Ser(Bzl) | L-hSer | L-Thr(Bzl) |
|---|---|---|---|---|---|---|---|---|---|---|---|---|---|---|---|---|---|---|---|---|
| WT | 0 | 0 | 0 | 0 | 0 | 0 | 0 | 0 | 0 | 0 | 0 | 0 | 5 | 0 | 100 | 0 | 45 | 8 | 0 | 36 |
| E47K | 0 | 0 | 0 | 0 | 0 | 0 | 0 | 0 | 0 | 0 | 0 | 0 | 0 | 0 | 100 | 0 | 38 | 0 | 0 | 36 |
| E47N | 0 | 0 | 0 | 0 | 0 | 0 | 0 | 0 | 0 | 0 | 0 | 0 | 0 | 0 | 100 | 0 | 50 | 16 | 17 | 45 |
| L50F | 0 | 0 | 0 | 0 | 0 | 0 | 0 | 0 | 0 | 0 | 0 | 0 | 0 | 0 | 100 | 0 | 0 | 0 | 0 | 0 |
| L89F | 0 | 0 | 0 | 0 | 0 | 0 | 0 | 0 | 0 | 0 | 0 | 0 | 0 | 0 | 100 | 0 | 41 | 0 | 0 | 45 |
| K90R | 0 | 0 | 0 | 0 | 0 | 0 | 0 | 0 | 0 | 0 | 0 | 0 | 0 | 0 | 100 | 0 | 34 | 0 | 0 | 30 |
| P108S | 0 | 0 | 0 | 0 | 0 | 0 | 0 | 0 | 0 | 0 | 0 | 0 | 0 | 0 | 100 | 0 | 33 | 0 | 0 | 56 |
| P132H | 0 | 0 | 0 | 0 | 0 | 0 | 0 | 0 | 0 | 0 | 0 | 0 | 0 | 0 | 100 | 0 | 49 | 0 | 0 | 30 |
| A260V | 0 | 0 | 0 | 0 | 0 | 0 | 0 | 0 | 0 | 0 | 0 | 0 | 0 | 0 | 100 | 0 | 28 | 0 | 0 | 17 |
| G283S | 0 | 0 | 0 | 0 | 0 | 0 | 0 | 0 | 0 | 0 | 0 | 0 | 0 | 0 | 100 | 0 | 35 | 0 | 0 | 29 |
| S284G | 0 | 0 | 0 | 0 | 0 | 0 | 0 | 0 | 0 | 0 | 0 | 0 | 0 | 0 | 100 | 0 | 33 | 0 | 0 | 25 |

| Variants | L-Cys(Bzl) | L-Cys(MeBzl) | L-Cys(4-MeOBzl) | L-Met | L-Met(O) | L-Met(O)$_2$ | L-Nle(O-Bzl) | L-Phg | L-hPhe | L-Chg | L-Cha | L-hCha | L-Igl | L-1-Nal | L-2-Nal | L-Bip | L-Bpa | L-2-Aoc | L-hLeu | L-Tle |
|---|---|---|---|---|---|---|---|---|---|---|---|---|---|---|---|---|---|---|---|---|
| WT | 0 | 0 | 15 | 0 | 0 | 0 | 0 | 0 | 0 | 0 | 0 | 0 | 0 | 0 | 0 | 0 | 0 | 0 | 0 | 50 |
| E47K | 0 | 0 | 14 | 0 | 0 | 0 | 0 | 0 | 0 | 0 | 0 | 0 | 0 | 0 | 0 | 0 | 0 | 0 | 0 | 35 |
| E47N | 0 | 0 | 27 | 14 | 0 | 0 | 0 | 0 | 0 | 0 | 0 | 0 | 0 | 10 | 0 | 0 | 0 | 0 | 0 | 49 |
| L50F | 0 | 0 | 0 | 0 | 0 | 0 | 0 | 0 | 0 | 0 | 0 | 0 | 0 | 0 | 0 | 0 | 0 | 0 | 0 | 0 |
| L89F | 0 | 0 | 10 | 0 | 0 | 0 | 0 | 0 | 0 | 0 | 0 | 0 | 0 | 0 | 0 | 0 | 0 | 0 | 0 | 36 |
| K90R | 0 | 0 | 0 | 0 | 0 | 0 | 0 | 0 | 0 | 0 | 0 | 0 | 0 | 0 | 0 | 0 | 0 | 0 | 0 | 36 |
| P108S | 0 | 0 | 16 | 0 | 0 | 0 | 0 | 0 | 0 | 0 | 0 | 0 | 0 | 0 | 0 | 0 | 0 | 0 | 0 | 38 |
| P132H | 0 | 0 | 16 | 0 | 0 | 0 | 0 | 0 | 0 | 0 | 0 | 0 | 0 | 0 | 0 | 0 | 0 | 0 | 0 | 44 |
| A260V | 0 | 0 | 0 | 0 | 0 | 0 | 0 | 0 | 0 | 0 | 0 | 0 | 0 | 0 | 0 | 0 | 0 | 0 | 0 | 39 |
| G283S | 0 | 0 | 18 | 0 | 0 | 0 | 0 | 0 | 0 | 0 | 0 | 0 | 0 | 0 | 0 | 0 | 0 | 0 | 0 | 28 |
| S284G | 0 | 0 | 15 | 0 | 0 | 0 | 0 | 0 | 0 | 0 | 0 | 0 | 0 | 0 | 0 | 0 | 0 | 0 | 0 | 26 |

| Variant | L-Arg(NO$_2$) | L-Tyr(Me) | L-Tyr(2,6-Cl$_2$-Bzl) | L-Tyr(Bzl) | L-Tyr(2-Br-Z) | L-hTyr | L-hTyr(Me) | L-Nva | L-His(Bzl) | L-2-Abz | L-3-Abz | L-4-Abz | Inp | L-Gla | L-Phe(4-NO2) | Aib | L-hSer(Bzl) | TmbGly | L-Nle |
|---|---|---|---|---|---|---|---|---|---|---|---|---|---|---|---|---|---|---|---|
| WT | 0 | 0 | 0 | 0 | 0 | 0 | 0 | 22 | 0 | 69 | 81 | 25 | 0 | 0 | 0 | 0 | 15 | 0 | 0 |
| E47K | 0 | 0 | 0 | 0 | 0 | 0 | 0 | 25 | 0 | 50 | 69 | 0 | 0 | 0 | 0 | 0 | 0 | 15 | 0 |
| E47N | 0 | 0 | 0 | 0 | 0 | 0 | 0 | 35 | 0 | 63 | 74 | 33 | 0 | 0 | 0 | 0 | 22 | 31 | 0 |
| L50F | 0 | 0 | 0 | 0 | 0 | 0 | 0 | 0 | 0 | 0 | 65 | 0 | 0 | 0 | 0 | 0 | 0 | 0 | 0 |
| L89F | 0 | 0 | 0 | 0 | 0 | 0 | 0 | 17 | 0 | 58 | 74 | 17 | 0 | 0 | 0 | 0 | 0 | 0 | 0 |
| K90R | 0 | 0 | 0 | 0 | 0 | 0 | 0 | 13 | 0 | 46 | 72 | 0 | 0 | 0 | 0 | 0 | 0 | 0 | 0 |
| P108S | 0 | 0 | 0 | 0 | 0 | 0 | 0 | 0 | 0 | 55 | 72 | 26 | 0 | 0 | 0 | 0 | 0 | 0 | 0 |
| P132H | 0 | 0 | 0 | 0 | 0 | 0 | 0 | 17 | 0 | 47 | 64 | 17 | 0 | 0 | 0 | 0 | 0 | 28 | 0 |
| A260V | 0 | 0 | 0 | 0 | 0 | 0 | 0 | 0 | 0 | 52 | 78 | 0 | 0 | 0 | 0 | 0 | 0 | 0 | 0 |
| G283S | 0 | 0 | 0 | 0 | 0 | 0 | 0 | 0 | 0 | 48 | 80 | 0 | 0 | 0 | 0 | 0 | 0 | 19 | 0 |
| S284G | 0 | 0 | 0 | 0 | 0 | 0 | 0 | 8 | 0 | 41 | 68 | 0 | 0 | 0 | 0 | 0 | 0 | 0 | 0 |

**Figure S6 continued.** Substrate specificity profile of SARS-CoV-2 M$^{pro}$ presented as heat maps using substrate library with unnatural amino acids assessing the P4 position with Ac-P4-Mix-Mix-Gln-ACC.

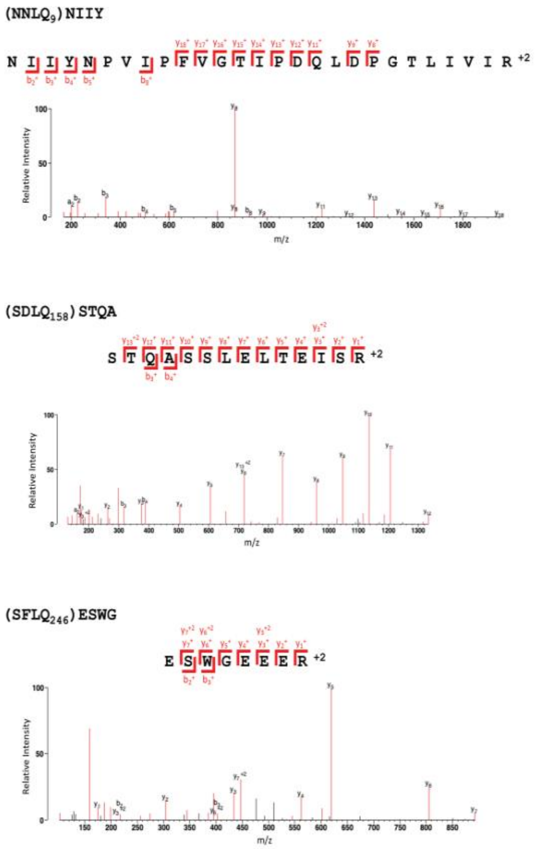


**Figure S7.** SARS-CoV-2 M$^{pro}$ cleaves Gal-8 at three sites as observed by mass spectrometry. Following incubation of Gal8 with SARS-CoV-2 Mpro for 2 h, the whole assay was analyzed by mass spectrometry. Three cleavage sites were identified for Gal-8 at $LQ_9$, $LQ_{158}$, and $LQ_{246}$.

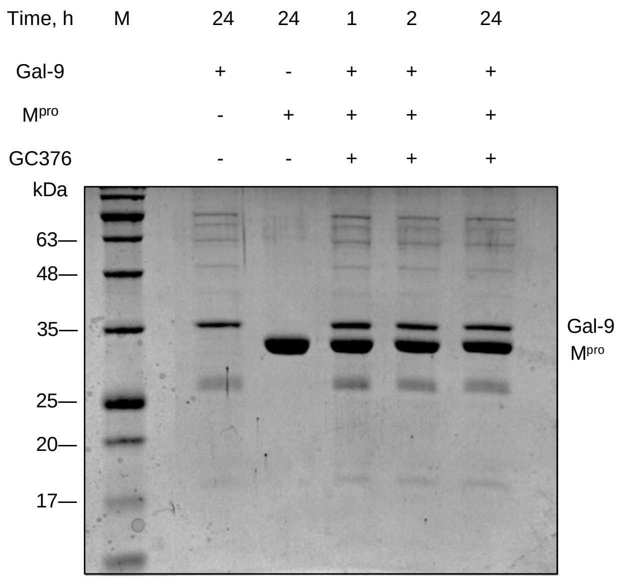


**Figure S8.** SDS-PAGE-based time-course of Gal-9 cleavage by SARS-CoV-2 M$^{pro}$. Gal-9 was incubated with the protease at 37°C and reaction was stopped at specific time points with M$^{pro}$ inhibitor, GC376.

A

B

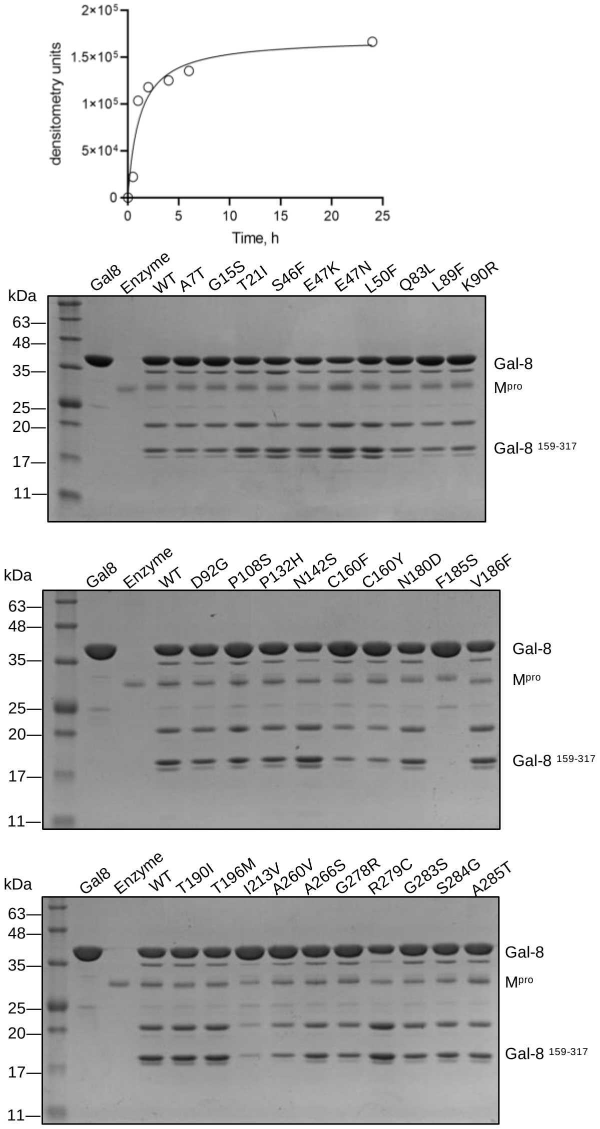


C

| *M^pro mutant* | *Gal8 ^159-317 amount, %* |
|---|---|
| *wild type* | 100 |
| *A7T* | 93.3 |
| *G15S* | 94.0 |
| *T21I* | 140.0 |
| *S46F* | 158.3 |
| *E47K* | 180.8 |
| *E47N* | 139.2 |
| *L50F* | 186.2 |
| *Q83L* | 105.3 |
| *L89F* | 81.9 |
| *K90R* | 104.0 |
| *D92G* | 81.8 |
| *P108S* | 84.2 |
| *P132H* | 89.2 |
| *N142S* | 120.2 |
| *C160F* | 48.6 |
| *C160Y* | 40.9 |
| *N180D* | 108.3 |
| *F185S* | 0 |
| *V186F* | 136.4 |
| *T190I* | 118.0 |
| *T196M* | 113.9 |
| *I213V* | 11.5 |
| *A260V* | 32.2 |
| *A266S* | 71.9 |
| *G278R* | 46.0 |
| *R279C* | 183.0 |
| *G283S* | 83.1 |
| *S284G* | 78.1 |
| *A285T* | 49.9 |

**Figure S9.** The cleavage of host cell substrate Gal-8 by SARS-CoV-2 $M^{pro}$ wild type and the mutants. (A) Densitometry analysis of dependence of Gal-8 $^{159-317}$ cleavage product formed on time upon SARS-CoV-2 $M^{pro}$ proteolysis. (B) Gal-8 was incubated with the proteases at 37°C for 2 hours, the reaction was stopped with $M^{pro}$ inhibitor, GC376 and SDS-PAGE-based densitometry analysis of Gal-8 $^{159-317}$ cleavage product was performed. (C) The results are depicted as a percent of generated Gal-8 $^{159-317}$ product in comparison to Gal-8 cleavage by the wild type.

**Table S1. The prevalence of mutations throughout the VOCs calculated as a percentage a certain mutation of all mutations in *nsp5* in a particular variant.** Only 'high coverage" sequences were considered at GISAID database (https://gisaid.org).

| | *Frequency of occurrence,%* | | | | |
|---|---|---|---|---|---|
| *Mutation* | *Alpha B.1.1.7* | *Beta B.1.351* | *Gamma P.1* | *Delta B.1.617.2* | *Omicron B.1.1.529* |
| *A7T* | 0.100 | 0.004 | 0.062 | 0.321 | 0.010 |
| *G15S* | 0.047 | 0.035 | 0.025 | 0.322 | 0.011 |
| *T21I* | 1.493 | 0.164 | 9.647 | 0.966 | 0.082 |
| *S46F* | 0.235 | 0.004 | 0.111 | 0.750 | 0.006 |
| *E47N* | 8.713 | 0.000 | 0.000 | 0.000 | 0.000 |
| *E47K* | 0.065 | 0.008 | 0.050 | 0.015 | 0.003 |
| *L50F* | 0.381 | 0.093 | 0.421 | 0.833 | 0.011 |
| *Q83L* | 0.005 | 0.000 | 0.000 | 0.016 | 0.003 |
| *L89F* | 0.720 | 0.008 | 2.093 | 0.307 | 0.034 |
| *K90R* | 23.672 | 99.887 | 16.223 | 21.764 | 0.581 |
| *D92G* | 0.320 | 0.000 | 0.025 | 0.184 | 0.003 |
| *P108S* | 5.379 | 0.277 | 2.885 | 1.873 | 0.056 |
| *P132H* | 0.011 | 0.000 | 0.000 | 0.131 | 99.947 |
| *N142S* | 0.000 | 0.000 | 0.000 | 0.007 | 0.000 |
| *C160F* | 4.109 | 0.035 | 0.693 | 0.086 | 0.003 |
| *C160Y* | 0.165 | 0.004 | 0.136 | 0.126 | 0.001 |
| *N180D* | 0.023 | 0.000 | 0.000 | 0.049 | 0.004 |
| *P184S* | 0.652 | 0.008 | 0.186 | 0.544 | 0.026 |
| *F185S* | 0.000 | 0.000 | 0.000 | 0.002 | 0.000 |
| *V186F* | 0.167 | 0.012 | 7.207 | 0.173 | 0.002 |
| *T190I* | 0.269 | 0.008 | 0.111 | 0.284 | 0.004 |
| *T196M* | 0.151 | 0.000 | 0.025 | 0.879 | 0.010 |
| *I213V* | 0.070 | 0.000 | 0.037 | 2.381 | 0.004 |
| *A260V* | 3.784 | 0.051 | 0.793 | 5.038 | 0.020 |
| *A266S* | 0.005 | 0.000 | 0.000 | 0.130 | 0.000 |
| *G278R* | 0.023 | 0.000 | 0.000 | 0.041 | 0.001 |
| *R279C* | 1.207 | 0.012 | 0.644 | 0.307 | 0.016 |
| *G283S* | 0.200 | 0.000 | 0.062 | 0.064 | 0.006 |
| *S284G* | 0.090 | 0.004 | 0.037 | 0.242 | 0.070 |
| *A285T* | 0.042 | 0.000 | 0.916 | 0.020 | 0.021 |
| *K90R P132H* | 0.000 | 0.000 | 0.000 | 0.004 | 0.786 |

**Table S2. Protein crystallography data collection and refinement statistics (molecular replacement).** Values in parentheses are for highest-resolution shell. Each data set was collected from a single crystal.

| | SARS-CoV-2 $M^{pro}$ A7T | SARS-CoV-2 $M^{pro}$ E47N | SARS-CoV-2 $M^{pro}$ E47K | SARS-CoV-2 $M^{pro}$ L50F |
|---|---|---|---|---|
| PDB Entry | 8DJJ | 8EJ9 | 8EJ7 | 8DKZ |
| **Data collection** | | | | |
| Space group | C 1 2 1 | C 1 2 1 | P 1 $2_1$ 1 | C 1 2 1 |
| Cell dimensions | | | | |
| *a, b, c* (Å) | 114.11 53.93 44.71 | 114.62 54.5 44.85 | 44.87 54.1 114.01 | 121.99 82.25 63.88 |
| a, b, g (°) | 90 102.073 90 | 90 101.186 90 | 90 101.028 90 | 90 90.673 90 |
| Resolution (Å) | 48.56 – 2.51 (2.6 – 2.51) | 28.11 – 2.5 (2.589 – 2.5) | 31.39 – 2.3 (2.382 – 2.3) | 46.47 – 3.0 (3.107 – 3.0) |
| Observations | 55345 (2927) | 62638 (6539) | 156339 (16305) | 84480 (8708) |
| $R_{merge}$ | 0.09358 (0.4553) | 0.135 (0.945) | 0.07774 (0.4827) | 0.1164 (1.088) |
| I / s I | 14.96 (2.99) | 8.73 (1.75) | 17.05 (4.18) | 13.15 (1.68) |
| Completeness (%) | 97.14 (80.04) | 97.00 (99.46) | 97.08 (98.35) | 94.59 (97.41) |
| Redundancy | 6.2 (4.0) | 6.7 (7.0) | 6.6 (6.8) | 6.7 (7.0) |
| CC1/2 | 0.997 (0.887) | 0.996 (0.785) | 0.999 (0.95) | 0.998 (0.598) |
| **Refinement** | | | | |
| Resolution (Å) | 48.56 – 2.51 | 28.11 – 2.5 | 31.39 – 2.3 | 46.47 – 3.0 |
| No. reflections | 8968 | 9246 | 23448 | 12109 |
| $R_{work}/R_{free}$ | 21.80/27.99 | 25.52/32.98 | 20.15/27.04 | 23.45/31.63 |
| Number of non-hydrogen atoms | 2407 | 2391 | 4960 | 4747 |
| Protein | 2372 | 2366 | 4756 | 4746 |
| Water | 35 | 25 | 204 | 1 |
| *B*-factors | | | | |
| Protein | 58.32 | 54.20 | 37.22 | 97.69 |
| Water | 48.13 | 50.15 | 36.75 | 52.84 |
| R.m.s. deviations | | | | |
| Bond lengths (Å) | 0.002 | 0.005 | 0.009 | 0.011 |
| Bond angles (°) | 0.53 | 0.69 | 1.05 | 1.42 |

**Table S3 continued. Protein crystallography data collection and refinement statistics (molecular replacement).** Values in parentheses are for highest-resolution shell. Each data set was collected from a single crystal.

| | SARS-CoV-2 $M^{pro}$ L89F | SARS-CoV-2 $M^{pro}$ K90R | SARS-CoV-2 $M^{pro}$ P132H | SARS-CoV-2 $M^{pro}$ T190I |
|---|---|---|---|---|
| PDB Entry | 8DKL | 8DKJ | 8DI3 | 8DK8 |
| **Data collection** | | | | |
| Space group | P 1 $2_1$ 1 | C 1 2 1 | C 1 2 1 | P 1 |
| Cell dimensions | | | | |
| *a, b, c* (Å) | 44.96 53.8 114.76 | 115.56 54.06 45.13 | 113.49 52.92 44.85 | 44.62 54.47 62.81 |
| a , b, g (°) | 90 101.355 90 | 90 101.079 90 | 90 102.897 90 | 115.719 99.579 90.061 |
| Resolution (Å) | 31.82 – 1.9 (1.968 – 1.9) | 27.5 – 2.11 (2.185 – 2.11) | 31.09 – 1.5 (1.554 – 1.5) | 43.85 – 2.6 (2.693 – 2.6) |
| Observations | 228777 (21074) | 107660 (10613) | 219276 (20962) | 52297 (4696) |
| $R_{merge}$ | 0.1039 (1.189) | 0.1519 (1.65) | 0.09456 (1.271) | 0.09051 (0.5521) |
| *I* / s*I* | 9.02 (1.38) | 9.05 (1.16) | 11.03 (1.61) | 9.36 (2.03) |
| Completeness (%) | 98.00 (95.68) | 98.83 (99.30) | 98.01 (96.60) | 93.92 (92.38) |
| Redundancy | 3.2 (5.2) | 6.9 (6.8) | 5.4 (5.2) | 3.4 (3.3) |
| CC1/2 | 0.998 (0.577) | 0.997 (0.408) | 0.997 (0.6) | 0.995 (0.752) |
| **Refinement** | | | | |
| Resolution (Å) | 31.82 – 1.9 | 27.5 – 2.11 | 31.09 – 1.5 | 43.85 – 2.6 |
| No. reflections | 41830 | 15693 | 40784 | 15109 |
| $R_{work}/R_{free}$ | 19.98/24.15 | 19.44/24.69 | 18.41/21.80 | 19.73/28.34 |
| Number of non-hydrogen atoms | 5062 | 2423 | 2691 | 4827 |
| Protein | 4762 | 2369 | 2373 | 4758 |
| Water | 300 | 54 | 318 | 69 |
| *B*-factors | | | | |
| Protein | 36.55 | 49.20 | 27.14 | 56.81 |
| Water | 41.17 | 48.34 | 37.78 | 47.59 |
| R.m.s. deviations | | | | |
| Bond lengths (Å) | 0.009 | 0.008 | 0.006 | 0.009 |
| Bond angles (°) | 1.09 | 0.92 | 0.95 | 1.22 |

**Table S3 continued. Protein crystallography data collection and refinement statistics (molecular replacement).** Values in parentheses are for highest-resolution shell. Each data set was collected from a single crystal.

| | SARS-CoV-2 $M^{pro}$ A260V | SARS-CoV-2 $M^{pro}$ G283S | SARS-CoV-2 $M^{pro}$ S284G |
|---|---|---|---|
| PDB Entry | 8DKH | 8DKK | 8DMN |
| **Data collection** | | | |
| Space group | C 1 2 1 | C 1 2 1 | C 1 2 1 |
| Cell dimensions | | | |
| *a, b, c* (Å) | 115.1 53.79 44.78 | 114.74 54.49 44.88 | 114.29 54.05 44.78 |
| a , b, g (°) | 90 101.58 90 | 90 101.029 90 | 90 101.018 90 |
| Resolution (Å) | 28.19 – 1.95 (2.02 – 1.95) | 38.45 – 2.0 (2.071 – 2.0) | 30.75 – 2.3 (2.382 – 2.3) |
| Observations | 128214 (13358) | 126086 (12750) | 174492 (17087) |
| $R_{merge}$ | 0.0656 (0.4928) | 0.1421 (1.231) | 0.1555 (1.272) |
| I / sI | 16.56 (4.05) | 10.95 (1.55) | 17.40 (5.64) |
| Completeness (%) | 97.20 (97.93) | 98.82 (99.02) | 97.44 (91.65) |
| Redundancy | 6.6 (7.0) | 6.9 (7.0) | 14.7 (15.6) |
| CC1/2 | 0.998 (0.945) | 0.997 (0.771) | 0.997 (0.955) |
| **Refinement** | | | |
| Resolution (Å) | 28.19 – 1.95 | 38.45 – 2.0 | 30.75 – 2.3 |
| No. reflections | 19144 | 18319 | 11760 |
| $R_{work}/R_{free}$ | 24.67/28.79 | 20.38/24.97 | 21.11/27.13 |
| Number of non-hydrogen atoms | 2537 | 2504 | 2457 |
| Protein | 2369 | 2369 | 2376 |
| Water | 168 | 135 | 81 |
| *B*-factors | | | |
| Protein | 33.01 | 44.00 | 37.73 |
| Water | 34.91 | 44.02 | 37.61 |
| R.m.s. deviations | | | |
| Bond lengths (Å) | 0.003 | 0.008 | 0.002 |
| Bond angles (°) | 0.61 | 0.95 | 0.53 |
